# The Halo Library, a Tool for Rapid Identification of Ligand Binding Sites on Proteins Using Crystallographic Fragment Screening

**DOI:** 10.1101/2022.10.17.512577

**Authors:** Ashima Chopra, Joseph D. Bauman, Francesc X. Ruiz, Eddy Arnold

## Abstract

X-ray crystallographic fragment screening (XCFS) uses fragment-sized molecules (∼60 – 300 Da) to access binding sites on proteins that may be inaccessible to larger drug-like molecules (>300 Da). Previous studies from our lab and others have shown that fragments containing halogen atoms have a higher binding occurrence compared to non-halogenated fragments. Specifically, we showed that 4-halopyrazoles hold potential for predicting the likelihood of success of a XCFS campaign. Here, we designed the Halo Library containing 46 halogenated fragments (including the “universal fragment” 4-bromopyrazole). The basis of fragment selection was presence of (at least) one halogen atom, and binding to or inhibitory activity against (at least) two targets in literature. The library was screened against crystals of HIV-1 reverse transcriptase with drug rilpivirine, yielding an overall hit rate of 26%. Two new binding sites were discovered in addition to previously reported sites, and several hot spots were identified (*i*.*e*., sites with multiple fragment hits). This small library may thus provide a convenient tool for assessing feasibility of a target for XCFS, mapping hot spots and cryptic sites, as well as finding fragment binders that can be useful for developing drug leads.

## INTRODUCTION

X-ray crystallographic fragment screening (XCFS) employs fragment-sized organic compounds (∼60 – 300 Da) to screen against crystals of biological drug targets. X-ray diffraction analysis of these crystals provides atomic models of the bound fragments, facilitating structure-based drug discovery (SBDD). It has an advantage over other high-throughput screening approaches, since fragment-sized compounds can access binding sites on the protein which may be unavailable to larger drug-like molecules.^1-4^ Moreover, it allows for exploring a larger chemical space with a limited number of screening compounds. Compared to high-throughput screening hits, typical fragment binders have a higher ligand efficiency (LE), a measure of binding affinity per non-hydrogen atom in the compound.^5^ LE is an established predictor of success in downstream drug development applications.^6^

Since its inception around 25 years ago, fragment-based drug design (FBDD) has come a long way – six drugs derived from FBDD have been approved, and multiple others are currently in clinical trials.^7, 8^ Having X-ray crystallography as the readout has important benefits over fragment screening by other methods like nuclear magnetic resonance (NMR) and surface plasmon resonance, as it allows structural visualization of binding sites and exploration of nearby binding pockets for fragment development.^1, 3^ In the past decade, this method’s relatively low throughput has been largely addressed through development of automated data collection and processing pipelines in synchrotrons and dedicated XCFS facilities such as XChem, FragMAX, Helmholtz-Zentrum Berlin (HZB), EMBL High Throughput Crystallization and Fragment Screening Facility (HTX Lab), and MASSIF-1.^9-13^

Fragments containing one or more halogen atoms have been shown to have a higher binding frequently against protein targets compared to non-halogenated fragments.^14-18^ Having more hits may provide alternative options for lead development in case a particular fragment development proves unconstructive. Halogen bonds likely contribute to their higher binding probability, employing the unique “σ-holes” used by halogens to strengthen contacts with side-chain oxygen and nitrogen atoms in the protein.^19, 20^ Presence of heavier halogens like bromine and iodine in the fragments makes them exceptionally well-suited for detection by X-ray crystallography due to their larger anomalous scattering compared to the carbon, oxygen, nitrogen, and sulfur atoms naturally occurring in proteins. This helps distinguish halogen electron density peaks and detect weak binding spots.^21^ Anomalous diffraction afforded by bromine and iodine can also help with phasing using single-wavelength or multi-wavelength anomalous dispersion methods, in cases where a good molecular replacement model is not available.^19, 22^ Last but not least, halogens have favorable chemical reactivity because they are good “leaving groups” in organic reactions. This facilitates fragment-to-lead development by employing halogen sites as reaction sites for chemical synthesis of lead compounds.^23, 24^ Nonetheless, the chemical reactivity of halogens can sometimes complicate chemical synthesis reactions in hit-to-lead optimization, and the potential of nonspecific interactions with hydrophobic sites in proteins has to be accounted for.^18^

Keeping the many advantages of halogenated fragments in mind, we developed the Halo Library for XCFS. It consists of 46 fragments containing one or more fluorine, chlorine, bromine, or iodine atoms. The library includes 32 fragment hits or analogs (differing from a fragment hit by the positioning or presence of one substituent) from our in-house fragment library which has been tested against HIV-1 reverse transcriptase (RT), influenza endonuclease, and integrase.^14, 25, 26^ 14 additional fragments were chosen that either show inhibition activity against at least two targets in literature, bind to at least two protein targets in the Protein Data Bank (PDB), or have analogs with the aforementioned properties (checked at https://pubchem.ncbi.nlm.nih.gov/). Proof-of-concept testing for the Halo Library was carried out against HIV-1 RT crystals in complex with the non-nucleoside drug rilpivirine (RPV).^27^ The results of a fragment screening campaign against these crystals have been previously published by our lab, where a 775-fragment library was soaked as cocktails into crystals of HIV-1 RT with RPV, laying the foundation for XCFS with the crystal system used in the current study.^14^

Anti-retroviral therapy (ART) for HIV-1 suffers from selection and accumulation of drug-resistance mutations, necessitating the development of new drugs.^28^ The enzyme RT is an exceptionally valuable drug target for HIV, because of its essential role in reverse transcribing the RNA genome into DNA during viral replication.^29^ About half of the drugs currently used in ART target the RT enzyme, highlighting its desirability as a drug target.^30^ Current ART regimens generally include at least two, and up to three RT inhibitors.^31, 32^ Presently used drugs targeting HIV-1 RT are primarily divided into: 1) nucleoside reverse transcriptase inhibitors (NRTIs), which act as chain terminators; and 2) non-nucleoside reverse transcriptase inhibitors (NNRTIs) – which act as allosteric inhibitors by altering RT conformation and disrupting its function. In addition to NRTIs and NNRTIs, novel mechanisms of HIV-1 RT inhibition have opened fresh avenues for SBDD against RT.^32, 33^

We report here the Halo Library screening of HIV-1 RT crystals bound to the NNRTI drug RPV, with an overall hit rate of 26% (vs. 4.4% for 775 fragments screened in 2013), yielding 12 binders from the 46 compounds tested. Two binding sites on HIV-1 RT previously unknown in literature were found to bind fragments. Multiple fragments were found to bind previously established inhibitory binding pockets including the Knuckles and NNRTI Adjacent site. Three fragments inhibit RT polymerization in the high millimolar range. This sets the foundation for fragment development in the inhibitory pockets by fragment merging, linking, and growing. Notably, the Halo Library can be used as a small-scale tool for rapidly and efficiently predicting likelihood of success of a larger XCFS campaign for a given crystal system. In addition, Halo Library screening holds promise as means to discover novel druggable sites on proteins that have not been reported in the literature.

## RESULTS AND DISCUSSION

### Library design

Of the 46 compounds in the Halo Library, 32 were halogenated fragments picked from our in-house library that was used for screening HIV-1 RT, influenza endonuclease, and integrase.^14, 25, 26^. Out of the 32, seven were pre-determined RT-RPV binders from our 2013 campaign.^14^ 14 fragments were purchased in addition. All fragments chosen showed inhibition activity against, or binding to, at least two targets in literature; or had chemical analogs that did the same. This strategy is hypothesized to yield a library that will have high binding occurrences in future XCFS campaigns as well.

It is worthwhile to note that in the previous XCFS RT campaign, soaking was done in cocktails of 4 – 8 fragments at 20 mM, which may have precluded identification of weakly bound fragments.^14^ However, for the Halo Library screening, soaks were done with individual fragments at 20 mM, which may also reduce chances of inter-fragment reactivity or competition for binding sites.

### Crystal growth and fragment soaking

RT52A, an RT construct engineered to produce high-resolution crystals, was used for this campaign.^27^ It is an enzymatically active and non-drug resistant mutant of RT. Purified protein was incubated and co-crystallized with RPV.^14, 27^ RPV binds in the NNRTI pocket of the enzyme, causing a conformational change.^34^ Co-crystallizing with RPV leads to locking of the enzyme in this conformation and prevents crystal damage caused by fragment binding in the NNRTI pocket. The co-crystals diffract reproducibly between 1.8 – 2.2 Å resolution, are very robust, and tolerant towards fragment soaking.

Fragment soaking was performed by harvesting RT-RPV crystals into crystal well solution mixed with cryoprotectants and fragment stock solutions (in dimethyl sulfoxide, DMSO) for two hours. Presence of 20% (v/v) DMSO and 5% ethylene glycol in the soaking solution allowed for efficient cryoprotection. The solution also contained 80 mM L-arginine which assists fragment dissolution. This is suspected to be due to the π-π stacking between the guanidinium group of arginine and aromatic rings on fragments, which helps counter the hydrophobic nature of many fragments, thereby facilitating solubilization.^14^ Most fragments can therefore be readily screened at a concentration of 20 mM.

A DMSO-only dataset was collected as a “blank” for creation of an isomorphous difference map *i*.*e*., F_obs(fragment soaked)_ – F_obs(DMSO blank)_ (*F*_o_ – *F*_o_) map. This helped in rapidly identifying electron density changes (visualized at 3σ) that are indicative of fragment binding. Ligand fitting was followed by complete data refinement and preparation of the coordinate models for PDB submission. Data collection and refinement statistics are presented in Table S2 in the Supporting Information. Difference electron density maps used for fitting the bound fragments are shown in Table S3.

### Halo Library screening overview

The 46 compounds of the Halo Library were tested over two rounds; the first round included 43 fragments which were available at the time of the synchrotron data collection, and the second round encompassed three additional fragments and unsuccessful datasets from the first round. Individual soaking was conducted with each fragment, and a total of 12 binders were found, giving a hit rate of 26% for this screening. Binding sites for all fragments are shown in Figure 1. Three of those binders were known from our previous campaign at the time of compiling the library. The hit rate excluding those fragments was found to be 20% (9 binders out of 43).

**Figure 1.**
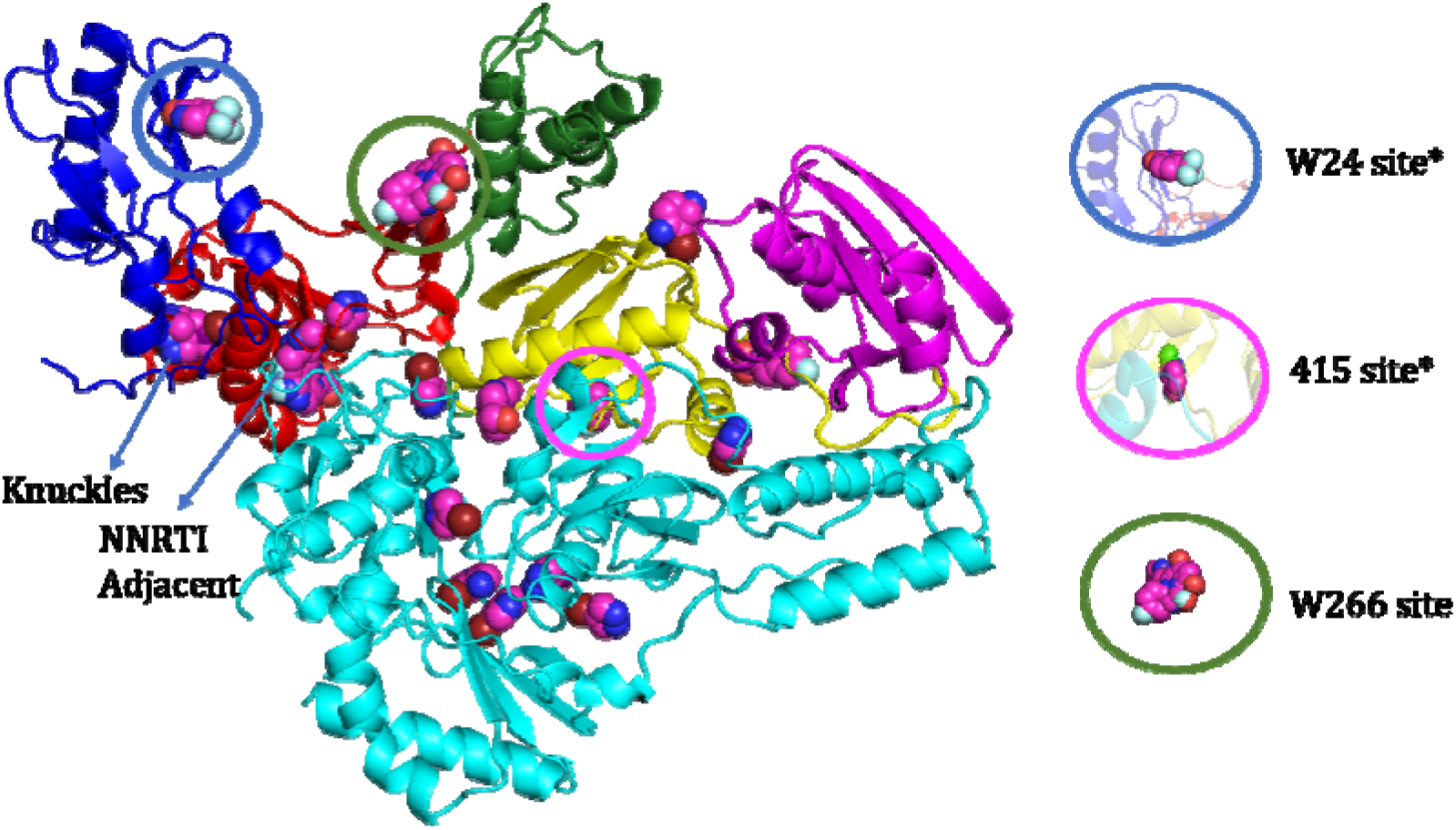
HIV-1 reverse transcriptase-rilpivirine (RT-RPV) with bound Halo Library fragments (magenta spheres). RT is color coded to show fingers (dark blue), palm (red), thumb (green), connection (yellow), and RNase H (magenta) subdomains of the p66 subunit; p51 subunit is shown in cyan. RPV is omitted for clarity. Ovals on the right show three novel RT binding sites; new sites discovered in this campaign are marked with an asterisk. Previously identified hotspots (Knuckles and NNRTI Adjacent site) are indicated.

The binders contained 7 fluorinated fragments (out of 24), 4 brominated fragments (out of 21), and 1 chlorinated and iodinated fragment each (out of 1 for each). This yields a hit rate of 29% for the fluorinated fragments and 19% for the brominated fragments (similar to the 24.1% and 23.5% hit rate respectively, in our 2013 campaign)^14^, which can be explained using unique interactions afforded by these two halogens. Fluorine, being the most electronegative element in the periodic table, is capable of short multipolar interactions of the type C-F…C=O in which fluorine interacts with the carbon in a carbonyl group.^28^ It is also a good hydrogen bond acceptor, and can form short contacts to hydrogen bond donors. In addition, the C-F bond is unique in its strength and hydrophobic nature which may help the fluorinated fragments access hydrophobic pockets on proteins.^35^ These factors may contribute to the higher hit rate of fluorinated fragments. Additionally, the favorable hit rate of Halo Library fluorinated fragments suggests the applicability of this subset of fragments for orthogonal fragment screening with ^19^F-NMR.^36^ On the other hand, bromine shows high halogen bonding tendencies due to the polarizability of its larger electron cloud. This leads to diffuse electronic orbitals and σ-holes which make attractive electrostatic interactions with carbonyl oxygen atoms.^37^

Six sites not seen in the 2013 study were found in this campaign. A comparison of the fragment-binding sites from the previous campaign and Halo Library screening is shown in Figure S1. ^14^ To assist in evaluating the relationship between these sites and the sites previously known in literature, we used the ligand pose comparison SIENA module in the proteins.plus server^38^, showing that two of those sites are previously unknown.^14^ The W24 site and 415 site, in addition to other potentially druggable sites found to bind fragments in this campaign, are discussed below.

### RT polymerization activity assay

A colorimetric assay (Roche), was used to determine the effect of fragment binders on RT polymerization activity.^39^ The widely utilized PicoGreen assay ^40^ was not effective for assessing fragment inhibition, as unspecific quenching of PicoGreen fluorescence was caused by several Halo Library fragments (data not shown). The assay used follows a sandwich ELISA protocol that leads to the formation of a colorimetric product, causing an absorbance increase which is directly proportional to RT activity. The absorbance values for RT activity with fragment and with vehicle [5% DMSO (v/v)] were determined, and percent activity was calculated. These values are shown in Table 1. Eight fragments inhibited RT activity by greater than fifty percent at 5 mM. Out of these, **HL6** (binding in the NNRTI Adjacent site, 428 site, and two p51 sites)^14^, **HL15** (binding in the 415 site) and **HL20** (binding in the NNRTI Adjacent site) were found to show >99% inhibition, and were investigated for IC_50_ and LE values (Table 1 and Supporting Information Figure S3). **HL6** shows an IC_50_ of 2.1 mM, with a favorable LE value of 0.47 (for reference, fragment-like leads typically should have at least a ligand efficiency of 0.38).^6^ It is yet to be discerned if the inhibitory activity is due to binding at the NNRTI Adjacent site, 428 site, or a combination of both. It is possible that binding at the p51 sites also contributes to inhibitory activity. **HL15** has an IC_50_ value of 2.1 mM, with LE value of 0.36. This presents a promising avenue for drug development at this newly discovered site, by fragment growth (see below). The NNRTI Adjacent binder **HL20** shows an IC_50_ of 2.9 mM with an LE of 0.28, presenting a similar scaffold to the 2013 NNRTI Adjacent binder (PDB ID 4KFB).^14^

**Table 1.**
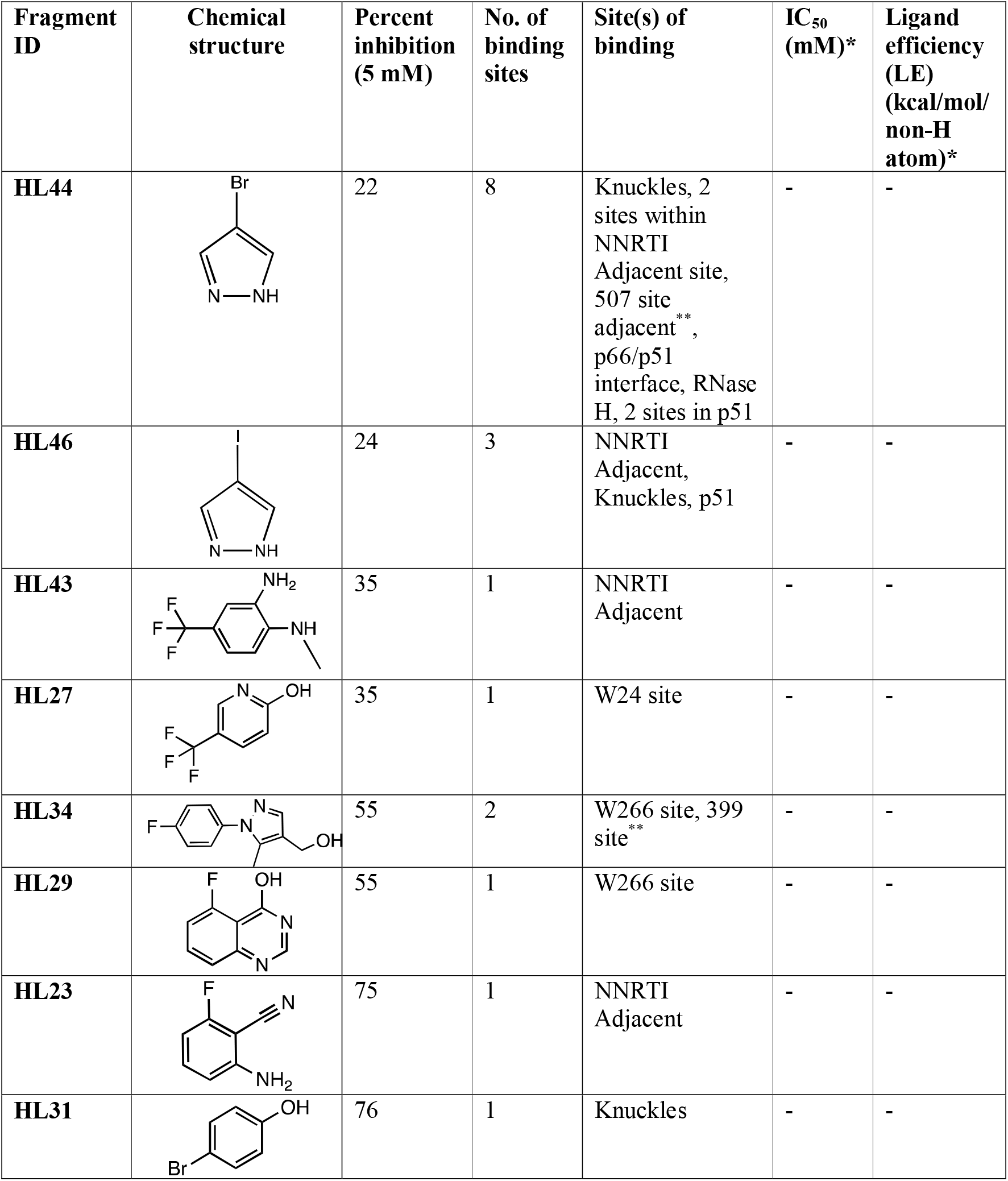
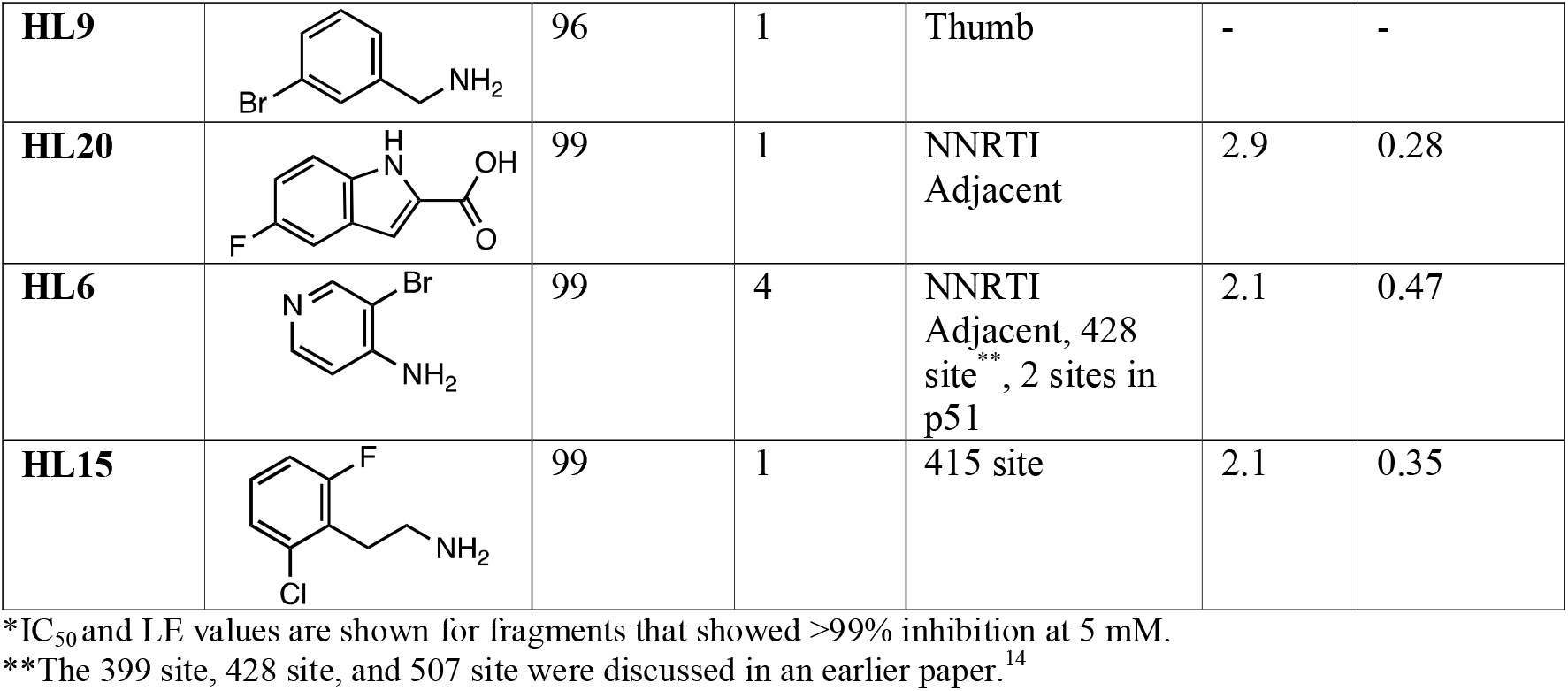
Inhibition of RT activity upon Halo Library fragment binding shown as a percentage compared to apo RT.

| Fragment ID | Chemical structure | Percent inhibition (5 mM) | No. of binding sites | Site(s) of binding | $IC_{50}$ (mM)* | Ligand efficiency (LE) (kcal/mol/non-H atom)* |
| --- | --- | --- | --- | --- | --- | --- |
| **HL44** |  | 22 | 8 | Knuckles, 2 sites within NNRTI Adjacent site, 507 site adjacent **, p66/p51 interface, RNase H, 2 sites in p51 | - | - |
| **HL46** |  | 24 | 3 | NNRTI Adjacent, Knuckles, p51 | - | - |
| **HL43** |  | 35 | 1 | NNRTI Adjacent | - | - |
| **HL27** |  | 35 | 1 | W24 site | - | - |
| **HL34** |  | 55 | 2 | W266 site, 399 site** | - | - |
| **HL29** |  | 55 | 1 | W266 site | - | - |
| **HL23** |  | 75 | 1 | NNRTI Adjacent | - | - |
| **HL31** |  | 76 | 1 | Knuckles | - | - |
| **HL9** |  | 96 | 1 | Thumb | - | - |
| **HL20** |  | 99 | 1 | NNRTI Adjacent | 2.9 | 0.28 |
| **HL6** |  | 99 | 4 | NNRTI Adjacent, 428 site **, 2 sites in p51 | 2.1 | 0.47 |
| **HL15** |  | 99 | 1 | 415 site | 2.1 | 0.35 |

\*IC<sub>50</sub> and LE values are shown for fragments that showed >99% inhibition at 5 mM.
\*\*The 399 site, 428 site, and 507 site were discussed in an earlier paper.<sup>14</sup>

### Fragment binding and development analysis

Figure 1 allows visualization of the overall picture of Halo Library screening with RT-RPV. The results of this campaign present opportunities for fragment development, especially at the sites found to bind more than one fragment. The fragment binders at these sites can be merged (*e*.*g*., at the NNRTI Adjacent site, W266 site, and Knuckles); or linked at sites where they bind close to a previously known ligand to form a larger binder (*e*.*g*., at the NNRTI Adjacent site). Binding sites of hits found to be inhibitory in our activity assay (Table 1) were assessed for their “druggability” to predict success in downstream applications using the DoGSiteScorer server (Table 2).^41, 42^ This tool helps predict the likelihood of a binding site to be a useful for drug binding by assessing its geometry and physico-chemical properties, which have previously proved useful to accurately predict the druggability of a target.^43-45^ Factors like volume of the pocket, enclosure within the protein, and hydrophobicity of residues in the pocket are calculated for target assessment. One drawback of this approach may be that it does not completely account for the solvent channels between nearby sub-pockets which can be exploited to create a binding site with larger volume and likely higher druggability score, as is discussed below for the NNRTI Adjacent and Knuckles sites. However, detailed examination of these solvent channels can help expand this analysis. Known NRTI and NNRTI drug target sites are shown as reference for the sake of comparison.

**Table 2.**
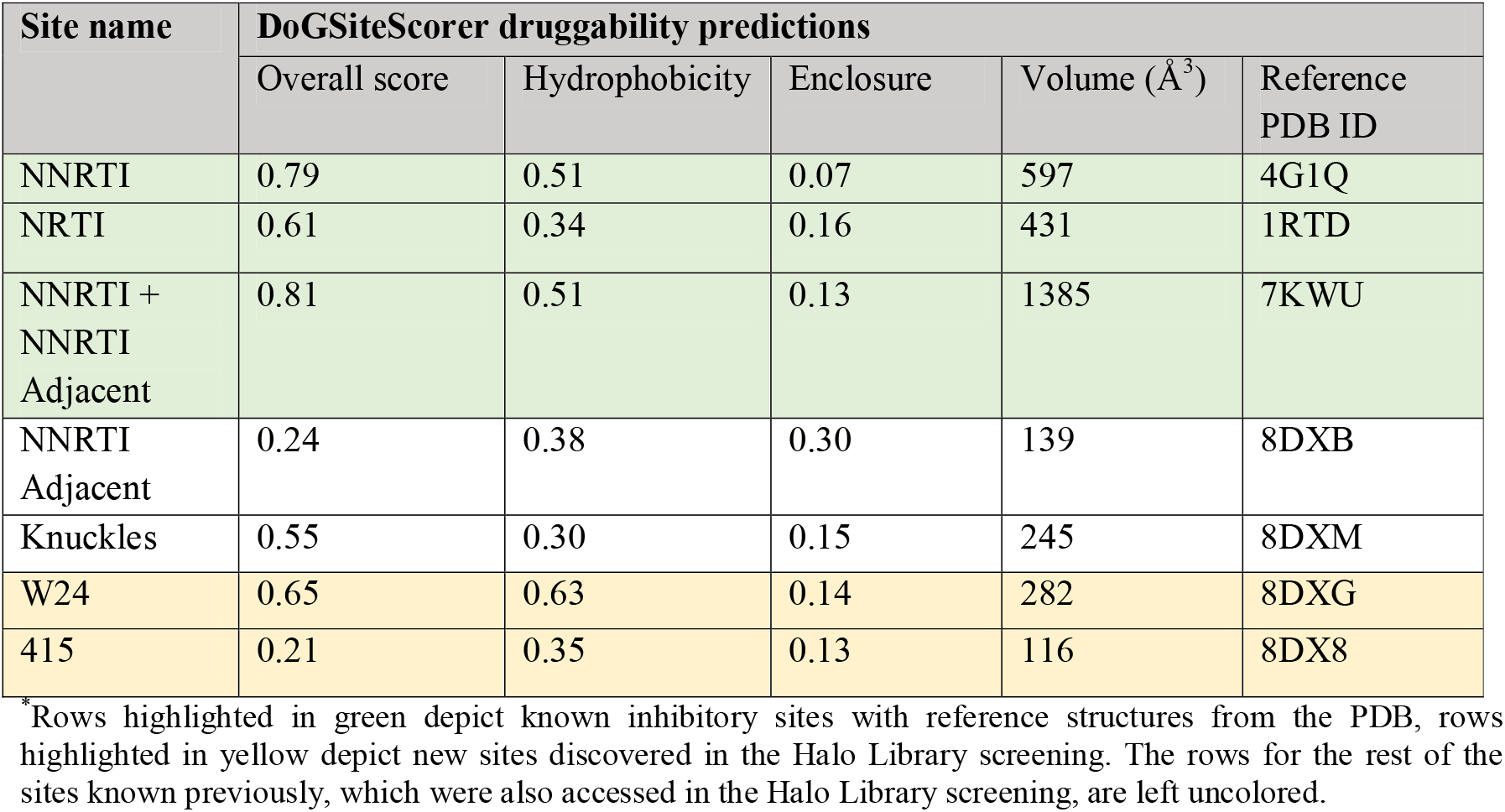
DoGSite druggability assessment for binding sites on HIV-1 RT^*^.^42^.

| Site name | DoGSiteScorer druggability predictions |  |  |  |  |
| --- | --- | --- | --- | --- | --- |
|  | Overall score | Hydrophobicity | Enclosure | Volume (Å<sup>3</sup>) | Reference PDB ID |
| NNRTI | 0.79 | 0.51 | 0.07 | 597 | 4G1Q |
| NRTI | 0.61 | 0.34 | 0.16 | 431 | 1RTD |
| NNRTI +<br>NNRTI<br>Adjacent | 0.81 | 0.51 | 0.13 | 1385 | 7KWU |
| NNRTI<br>Adjacent | 0.24 | 0.38 | 0.30 | 139 | 8DXB |
| Knuckles | 0.55 | 0.30 | 0.15 | 245 | 8DXM |
| W24 | 0.65 | 0.63 | 0.14 | 282 | 8DXG |
| 415 | 0.21 | 0.35 | 0.13 | 116 | 8DX8 |

<sup>\*</sup>Rows highlighted in green depict known inhibitory sites with reference structures from the PDB, rows highlighted in yellow depict new sites discovered in the Halo Library screening. The rows for the rest of the sites known previously, which were also accessed in the Halo Library screening, are left uncolored.

### NNRTI Adjacent site

The NNRTI Adjacent site, which is close to the entrance channel of the NNRTI-binding pocket (NNIBP), was found to bind six fragments from the Halo Library (Figure 2). All fragments were found to overlap to varying extents depending on the fragment size, underlining the importance of this site as a hotspot for probes or drugs (Figure 2G). Two of the fragments, **HL6** and **HL20**, inhibit RT polymerization with an IC_50_ of 2.1 and 3.0 mM respectively, and corresponding LE values of 0.47 and 0.28 kcal/mol/non-H atom (Table 1; although **HL6** binds multiple sites, and its inhibition activity cannot be attributed to binding at this site alone). Because of the NNRTI Adjacent site’s relatively small pocket volume, its druggability score is low at 0.24, even though the hydrophobicity value (0.38) is comparable to the NRTI site (0.34). Its enclosure value is greater (0.30) than the NRTI site (0.16) and NNRTI site (0.07), suggesting that it is a deeper pocket. Binders at this site can be linked with NNRTI binders to increase overall pocket volume, using a solvent channel between the two sites as shown before.^46, 47^ Such inhibitors are shown in literature to target both the NNIBP and NNRTI Adjacent sites, with <5 nM EC_50_ values and possessing activity against multiple resistant mutants of HIV-1 RT.^47^ Intriguingly, our laboratory has observed the presence of ordered water networks surrounding the NNIBP, and specifically that the ordered water network near the entrance channel is in a continuum with an ordered water network in the NNRTI Adjacent site (PDB ID: 4G1Q).^14, 48^ The fact that the binders in this site displace ordered water molecules (Figure 2g) could entail an entropic gain, with the potential to increase the binding affinity of NNRTIs targeting both the NNIBP and the NNRTI Adjacent site.

**Figure 2.**
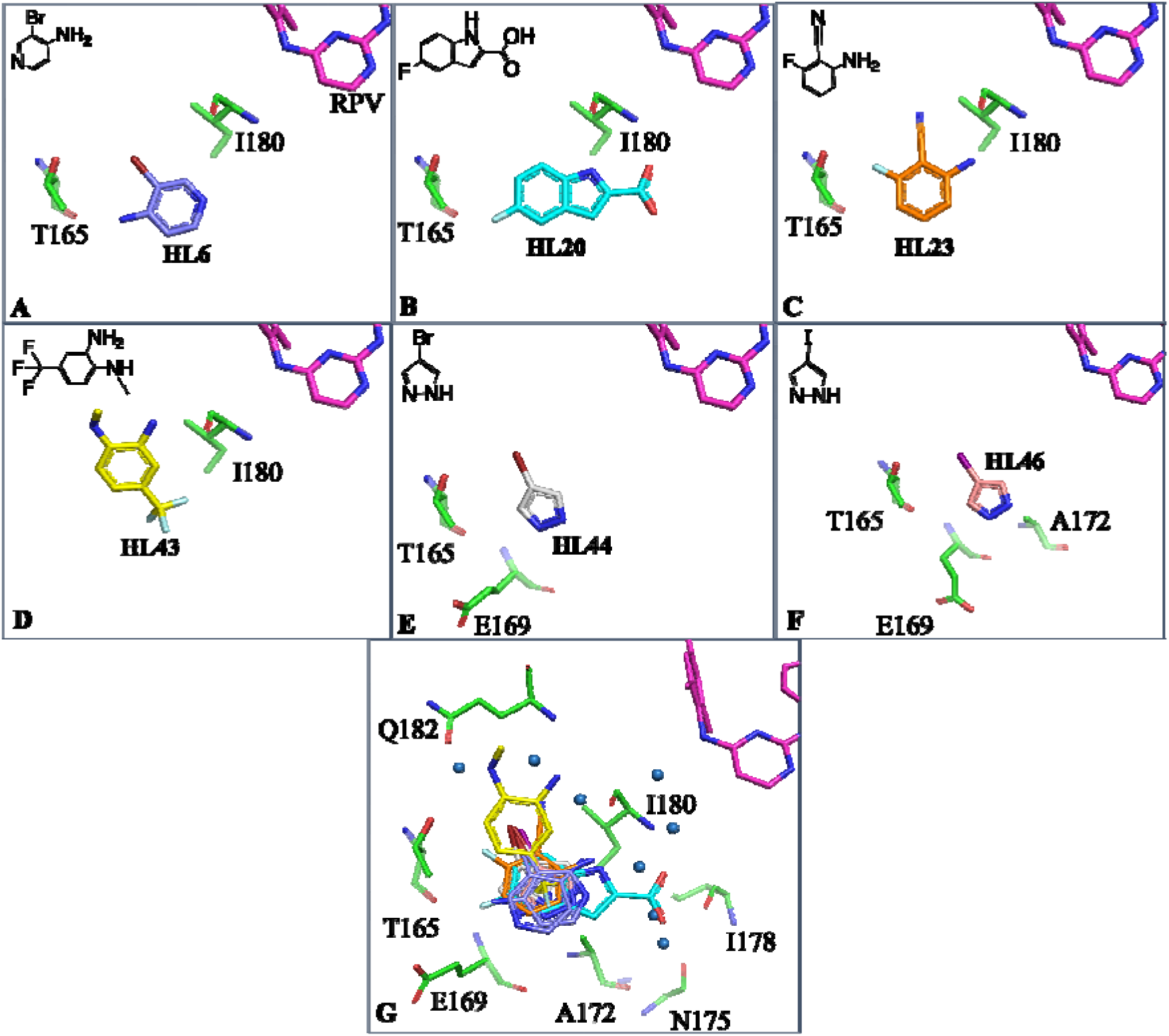
NNRTI Adjacent site. Panels (A-G) show Halo Library fragments as labeled in differently colored sticks. Directly interacting p66 residues are shown as green sticks. Rilpivirine is denoted as RPV in panel (A) and is shown in all panels for reference. Panel (G) shows all 6 fragments overlapping, with the water network as blue spheres, and p66 residues as green sticks which interact with fragments through one or more water-mediated hydrogen bonds. Water molecules depicted are corresponding to PDB ID 7KWU.

### W24 site

W24 lies in the fingers subdomain of p66 and undergoes substantial rearrangement to accommodate **HL27** in the newly discovered W24 site (Figure 3A). Side-chain rotation allows for π-π stacking of its indole ring with the aromatic ring of the fragment. Binding at this site is also facilitated by a short and presumably strong hydrogen bond (2.3 Å) between the aromatic ring nitrogen of the fragment and the main-chain carbonyl oxygen of K22, and a water-mediated bond with the side-chain oxygen of D76 (Figure 3B). This pocket has a druggability score of 0.65 and was found to be inhibitory for RT polymerase activity in our assay (Table 1). The high hydrophobicity value (0.63) likely contributes to the creditable druggability score, since the enclosure value (0.14), and volume of the pocket (282 Å^3^) are on the lower side for a drug-binding pocket (Table 2). A solvent channel close to **HL27** binding site offers an avenue for fragment growth which may help increase the volume of the pocket (Figure 3C).

**Figure 3.**
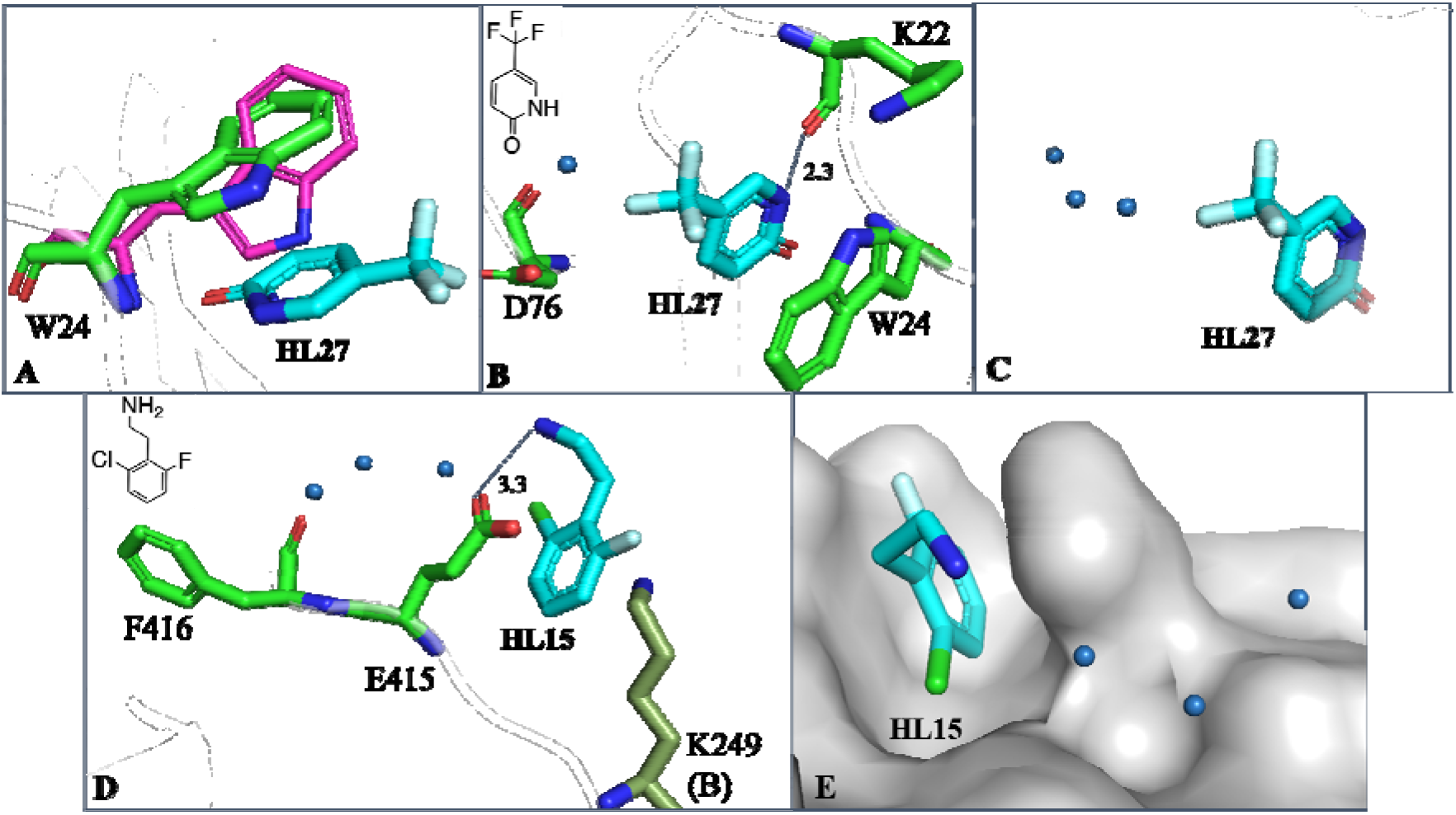
W24 site (A-C) and 415 site (D-E). Fragments are represented as cyan sticks and directly interacting p66 residues are shown as green sticks. (A) Shift of W24 upon fragment binding from magenta sticks to green sticks (compared to RT-RPV model, PDB ID 4G1Q). (B) W24 site showing **HL27** interacting with p66 residues. (C) Surface representation of p66 showing solvent channel for fragment expansion. (D) 415 site showing **HL15** interacting with p66 residues through ordered water molecules, and K249 from p51 (in dark green) of the symmetry mate. (E) Surface representation showing water channels for fragment expansion.

W24, alongside F61, constitutes an aromatic platform interacting with the nucleotide bases of the template strand overhang and is important for positioning of the template-primer substrate.^49, 50^ It is a well-conserved residue and occurs in the consensus sequence for RT across various clades (https://www.hiv.lanl.gov). Notably, mutation of W24 to a non-aromatic residue compromises dNTP incorporation, which makes the appearance of resistance mutations for this residue unlikely. ^49^ It is thus encouraging that **HL27** binding hinders polymerization activity (Table 1), as it likely has potential to be developed into a polymerization inhibitor which is less susceptible to drug resistant mutations.

### 415 site

This newly identified site lies in the connection subdomain of p66. The connection subdomain may be relevant for polypurine tract removal and RNase H cleavage specificity, and mutations in the connection have been described that cause resistance to RT inhibitors including NNRTIs.^51, 52^ This pocket binds **HL15** by hydrogen bonds between the amino group of the fragment and the side-chain oxygen of E415 in addition to an interaction with K249 (p51) in a neighboring RT molecule related by crystallographic symmetry (Figure 3D). Water molecules bridge interactions with F416. Additional hydrophobic interactions with K388 and K390 are not shown but likely contribute to the binding. This site utilizes an extensive water network to bind the fragment, which can be used for fragment extension (Figure 3D-E). **HL15** binding at this site inhibits RT activity with an IC_50_ of 2.1 mM and an LE of 0.35 (Table 1). Consistent with the literature,^53^ **HL15** may be inhibiting polymerization allosterically, although our data are insufficient to speculate by which mechanism.

### W266 site

W266 site binds two fragments, **HL29** and **HL34**, which exert stacking interactions with W266 in the base of thumb of the p66 subunit (Figure 4A-C). W266 is a part of the thumb subdomain that binds the nucleic acid on RT, being critical for its proper positioning.^54^ Hence, this site may be useful for drug development against HIV, as mutations are likely be detrimental for the virus. In addition, W266 is a highly conserved RT residue (https://www.hiv.lanl.gov) which is involved in interaction with the nucleic acid substrate, and its mutation blocks polymerization, RNase H and strand transfer activities.^55^ This could allow development of drugs at this site which may possess a high genetic barrier to resistance.

**Figure 4.**
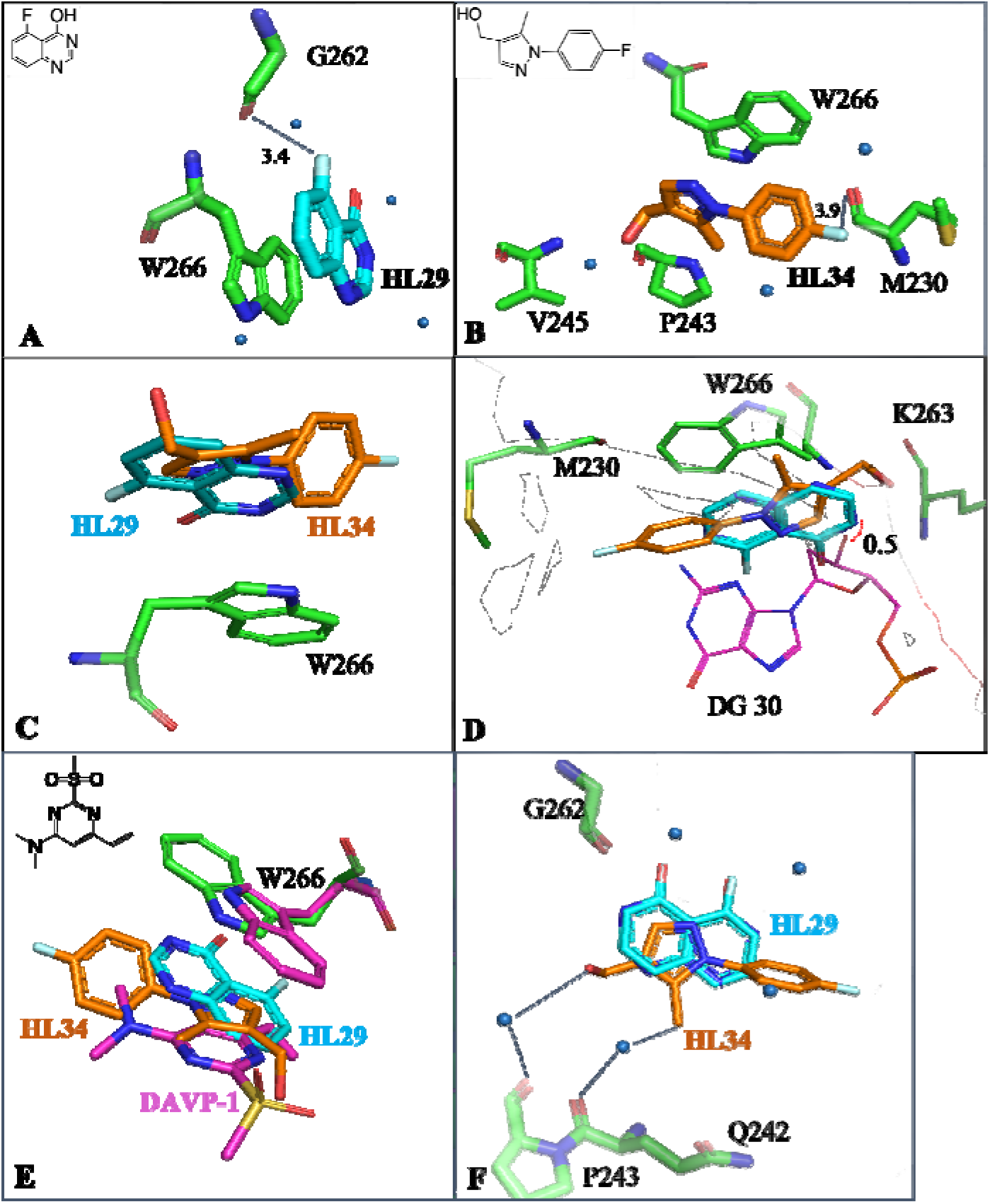
W266 site. Interacting p66 residues are shown as green sticks, **HL29** and **HL34** are shown as cyan and orange sticks respectively, water molecules are blue spheres. (A-B) Interaction of **HL29** and **HL34** with p66 residues and waters in the pocket. (C) Overlap of **HL29** and **HL34** stacking with the W266 side chain. (D) Overlay of fragment bound structures with nucleic acid bound RT (PDB ID 5D3G). **HL29** and **HL34** show small overlap with the sugar ring of primer strand nucleotide DG30 in 5D3G (magenta sticks). (E) Overlap of **HL29** and **HL34** with DAVP-1 (magenta sticks), with W266 in each structure (green sticks for **HL29** and **HL34** RT, magenta sticks for DAVP-1 RT) shown for reference. Structure of DAVP-1 is shown as an inset. (F) Water network in the W266 site for fragment extension.

**HL29** and **HL34** employ π-π stacking interactions with W266, and both show 55% inhibition of RT polymerization activity compared to unbound RT at 5 mM (Figure 4A-B and Table 1). **HL29** binds RT in two equivalent conformations (Figure 4A, one conformation is shown for clarity). In addition to stacking with W266, it forms a main-chain hydrogen bond with the carbonyl oxygen of G262, and additional hydrogen bonds with four water molecules. **HL34** shows water-bridged interactions with P243 and V245, and a hydrogen bond with the carbonyl oxygen of M230 (Figure 4B). The two fragments bind in this site with a significant overlap (Figure 4C).

When overlaid with a nucleic acid bound structure (PDB ID: 5D3G), the fragment-binding site partially overlaps with deoxyguanosine nucleotide 30 (DG 30), a position in the primer strand which is located three nucleotides downstream of the primer 3’ terminus. The substantial clash is likely significant enough to hamper nucleotide binding (Figure 4D). A previous two-stage fragment screening campaign (NMR and activity assays) detected a *p*-hydroxyaniline fragment acting as competitive inhibitor of the nucleic acid substrate, and low micromolar concentrations of the fragment inhibited HIV-1 replication in cell culture with a time-of-addition profile consistent with an RT inhibitor.^56^

The crystal structure of the inhibitor DAVP-1 [belonging to a class of 4-dimethylamino-6-vinylpyrimidine inhibitors, that act as nucleotide-competing (nc)RTIs] bound to RT showed binding in the same location, stacking against W266 (Figure 4E).^57-59^ However, these structures (in the absence of RPV) had the RT thumb in the “closed” conformation. Our structures with **HL29** and **HL34** capture the RT thumb in the “open” (or hyperextended) conformation, achieved by binding of NNRTI RPV. **HL29** and **HL34** could help to guide modifications of DAVP-1, by considering substituent modifications inspired by the binding locations and chemistry of the fragments. Potentially, they could act synergistically with RPV, while in its absence they may bind similarly to DAVP-1.

This site presents multiple opportunities for fragment development. The methyl group of **HL34** has a water-bridged interaction with P243 (similar to **HL29**), which can be used for fragment extension (Figure 4F). **HL34** is also in close contact with an entrapped water molecule that interacts with Q242, suggesting the replacement of this group by a hydrogen bond accepting group, *e*.*g*., trifluoromethyl (Figure 4F). Both mentioned water molecules are ordered, and conserved, as they were observed in the 1.51 Å resolution RT-RPV structure (PDB ID 4G1Q and Kuroda, D.G. *et al*., 2013).^60^ Three additional water molecules constitute an extensive water bonding network, bridging the fragments and residues G262, P243, and Q242, offers prospects for fragment development (Figure 4F). Drawbacks for this site may be that: i) it does not present a “pocket,” hence it could not be assigned a druggability score; and ii) binders here have to compete against the large nucleic acid substrate. This may be overcome by careful design of inhibitors that utilize a variety of π-π stacking, hydrogen bonding, and electrostatic interactions to overcome non-specific and/or weak binding.

### Knuckles site

This site lies between the fingers and palm subdomains of the p66 subunit (Figure 1), and was discussed in some detail earlier.^14^ In our previous campaign we found two fragment binders with inhibitory capability at the Knuckles site, out of which one was developed into a small molecule binder (PDB IDs 4IFY, 4IG3).^14^ In the current campaign, we found two Halo Library binders at this site. **HL31** binds using its hydroxyl and bromine substituents to form hydrogen bonds and halogen bonds respectively with main-chain carbonyl oxygen atoms of F160 and W212 (Figure 5A). The substituents are also in hydrogen bonding proximity of the main-chain amide nitrogen atom on M164. One water molecule in ∼4 Å proximity can be used as a directional guide for fragment expansion. This would help increase the volume of the druggable pocket from ∼250 Å^3^ to ∼400 Å^3^, including a 160 Å^3^ pocket that appears next to it in the DoGSiteScorer analysis (Figure S2). An example of fragment expansion at this site is demonstrated in PDB ID 4IG3. **HL46** (4-iodopyrazole) is the second binder at this site with its iodine atom overlapping with **HL31** (Figure 5C). The binding pocket consists of residues different from **HL31** (Figure 5B), including hydrogen bonds with the main-chain atoms of A114, S117, V118, and M164. It binds with partial occupancy which could explain its low inhibition – 24% at 5 mM compared to apo RT – as opposed to **HL31** which inhibits up to 75% RT activity at 5 mM (Table 1). The obtained hits confirm that the Knuckles site is a binding hot spot with inhibition capability.

**Figure 5.**
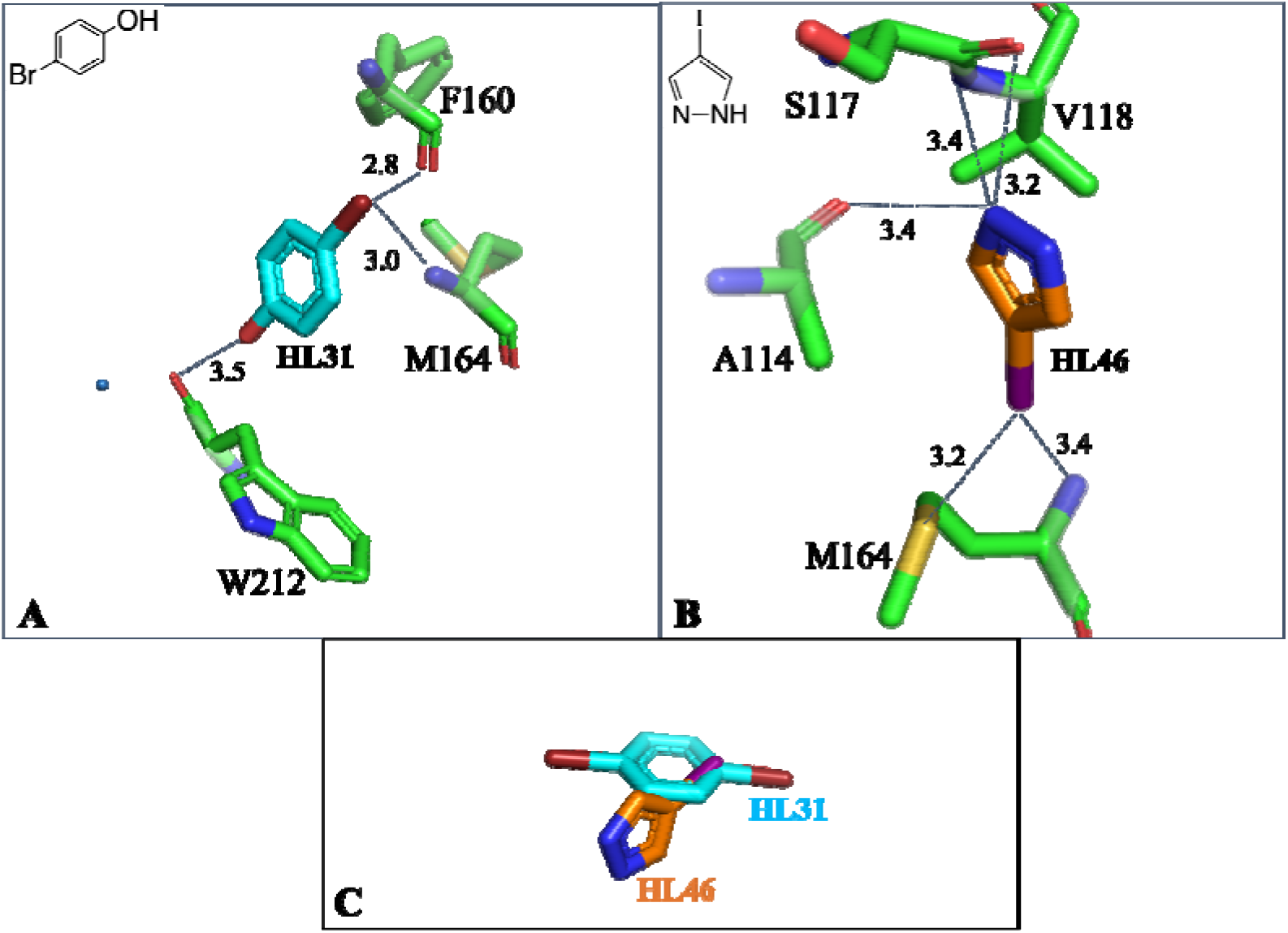
Knuckles site. (A) **HL31** (cyan sticks) and (B) **HL46** (orange sticks) interacting with p66 residues (green sticks). Significant hydrogen bonds are highlighted. (C) Overlap of the two fragments at the Knuckles site.

## CONCLUSION

In this work, we have designed a small fragment library, the Halo Library, based on the previous observations of unusually high hit rates of halogenated fragments in XCFS campaigns. As a proof-of-concept, we have applied XCFS using the Halo Library to HIV-1 RT, one of the most studied and screened protein targets.^32^ Nonetheless, Halo Library screening has unveiled new sites, confirmed binding hot spots, and provided meaningful avenues for further development. As in the previous campaign, we have performed an enzymatic activity assay with the obtained hits, to assess which binders bear more potential for development into inhibitors and/or probes.

The Halo Library may be an efficient and affordable way to assess feasibility of a target for XCFS, while probing ligandability, discovering new druggable sites, and providing paths for drug discovery. We are currently leveraging the Halo Library for novel drug targets, including the HIV-1 capsid protein. We have provided the chemical structures of all library members in the Supporting Information to allow independent testing of and experiments with the library.

## EXPERIMENTAL SECTION

### Fragment library preparation

14 out of the 46 fragments in the Halo Library were purchased from Sigma Aldrich and Alfa Aesar. 100 mM stock solutions of each were made by dissolving 2 mg of the fragment in an appropriate volume of DMSO. The remaining 32 fragments were chosen from our in-house fragment library. They were originally acquired from Maybridge, Sigma Aldrich, or Acros. 100 mM stocks in DMSO had been stored at - 80°C, and thawed before use. All fragments were obtained at >99% purity.

### Expression, purification, and crystallization

Construct RT52A of HIV-1 RT was expressed and purified as described previously.^27^ Purified protein at 20 mg/mL was prepared for crystallization by incubating it with RPV at 1:1.5 protein to drug molar ratio for 30 minutes at room temperature. Crystals were grown in 2 µL hanging drops at 4°C with 1:1 protein to well solution ratio. The well solutions contained 10 – 12% polyethylene glycol (PEG) 8,000, 4% PEG 400, 100 mM imidazole pH 6.4 – 6.5, 15 mM magnesium sulfate, 100 mM ammonium sulfate and 5 mM tris(2-carboxyethyl)phosphine. Pre-seeding was done using microseeds from previously generated and crushed RT52A-RPV crystals.

### Fragment soaking

Crystals of RT52A grown with RPV were harvested after two weeks and pre-screened for diffraction and DMSO tolerance using an in-house Rigaku R-Axis IV++. For pre-screening, crystals were swished through a cryo holding drop containing well solution, 80 mM L-arginine, 5% ethylene glycol and 20% DMSO. Data was collected overnight and checked to confirm protein diffraction. Soaking solutions were prepared by adding each fragment stock (100 mM) to a final concentration of 20 mM to a cryo solution containing well solution, 5% ethylene glycol, and 80 mM L-arginine. Soaking solution containing 20% DMSO instead of the fragment was used as the control. Approximately 50 – 60 RT-RPV crystals (for a 46-fragment campaign) were harvested into a 50 µL holding drop containing the well solution. This was followed by adding 50 µL of a solution containing well solution with 20% DMSO to the side of the holding drop, to achieve a 10% DMSO concentration in the holding drop and ease the crystals into the subsequent fragment-soaking solution. Crystals were then harvested into 1 µL of the soaking solution in a 96-well Intelli-Plate (Hampton Research) and the drops were sealed. They were incubated for 2 hours, followed by harvesting and flash-freezing in liquid nitrogen.

### Data collection and processing

Diffraction data was collected remotely at the 23-ID-D beamline of the Advanced Photon Source (APS) at Argonne National Laboratory (Lemont, IL), using pre-optimized collection settings. Crystal rastering was performed prior to data collection on the area of the crystal showing highest diffraction intensity (not including ice). Datasets were indexed, integrated and scaled using HKL2000.^61^ Rigid body refinement of scaled maps was performed in Phenix using 4G1Q as the reference model, to get a starting model and electron density maps. Resulting maps were used to make *F*_o_ – *F*_o_ maps, as well as polder OMIT maps.^62, 63^ *F*_o_ – *F*_o_ difference map peaks were visualized using Coot^64^ and checked for unmodeled blobs indicating fragment electron density. Fragment-bound crystal datasets were subjected to further processing, ligand fitting, and refinement cycles using Phenix.^62^ For datasets where *F*_o_ – *F*_o_ difference map density was incomplete, polder OMIT maps were used for fragment fitting. Figures showing protein-ligand interactions were made using PyMOL. Analysis of protein-fragment interactions was done manually in Coot, and aided by the PLIP server.^65^

### RT activity assay

Activity assays to see the inhibitory effect of Halo Library fragment binders on RT52A activity were performed using the Roche colorimetric assay kit purchased from Sigma Aldrich, using the absorbance measurement of 2,2_′_-Azino-bis(3-ethylbenzthiazoline-6-sulfonic acid) (ABTS) substrate at 405 nm (Sigma Aldrich Catalog # 11468120910).

Absorbance spectra of the fragments were measured between 200 – 900 nm wavelength to pre-screen for potential interference with ABTS absorbance. The λ_max_ for absorbance by the Halo Library binders was found to be in the range of 270-320 nm and showed no overlap with the λ_max_ for ABTS (405 nm).

All absorbance measurements were performed using SpectraMax iD3 multiplate reader (Molecular Dimensions). Stock solutions for the activity assay were prepared in diethyl pyrocarbonate-treated water using filter tips and RNase-free tubes to avoid RNase contamination during the assay. RT52A (30 nM) and inhibitor (5 mM Halo Library fragment) were pre-incubated at 37°C for one hour. Template-primer hybrid was incubated with the RT + inhibitor solutions for one hour. RT52A pre-incubated with RPV (10 µM) for 10 minutes at room temperature was used as a positive control for inhibition, and the enzyme with 5% DMSO was used as a negative control. Lysis buffer supplied with the kit, in the absence of any enzyme or inhibitor, was used as the blank. All concentrations mentioned are pre-assay concentrations before the components were mixed with each other. The DNA-RNA hybrid formed as a result of polymerization was then annealed to 96-well streptavidin plates and attached to anti-digoxigenin-peroxidase and horseradish peroxidase enzyme as described in the kit protocol. 200 µL ABTS substrate (1 tablet/ 5 mL substrate buffer) was incubated with the nucleic acid away from light for 20 minutes until a deep teal color was seen in the enzyme-only control, indicating appropriate substrate conversion into colorimetric product. The plates were then shaken for 30 s and absorbance was measured at 405 nm. Triplicate measurements were taken for each inhibitor-protein interaction, which were then averaged and normalized against the blank. Data were analyzed in MS Excel and percent inhibition values were calculated using absorbance values for RT52A incubated with Halo Library fragments, and RT52A with 5% DMSO.

IC_50_ values for fragments inhibiting RT activity >99% at 5 mM were calculated. Fragment concentrations were varied between 250 µM and 6 mM, and inhibition of RT52A was tested by measuring absorbance values of ABTS as described above. IC_50_ values were calculated using GraphPad Prism.

## SUPPORTING INFORMATION

Supporting Information provides additional figures to complement the discussion in text. Chemical names and SMILES strings of the Halo Library fragments are shared in Table S1.

### Accession Codes

Crystallographic datasets for the 12 RT-RPV Halo Library fragment binders were deposited to and are accessible from the PDB (https://www.rcsb.org/). PDB accession codes for all structures derived from this study are as follows: 8DX2, 8DX3, 8DX8, 8DXB, 8DXE, 8DXG, 8DXH, 8DXM, 8DXI, 8DXJ, 8DXK, and 8DXL and detailed in Supporting Information Table S2.

## ACKNOWLEDGMENTS

We are grateful to APS staff members for support and access to beamline 23-ID-D for data collection, and to all the members of the Arnold lab for helpful discussions. This work was supported by NIH grants U54 AI150472 (HIVE Center) and R01 AI026790 (both to E.A.). The authors declare no conflict of interest.

## ABBREVIATIONS USED

ABTS: 2,2′-Azino-bis(3-ethylbenzthiazoline-6-sulfonic acid)
APS: Advanced Photon Source
ART: antiretroviral therapy
DAVP-1: 4-dimethylamino-6-vinylpyrimidine
DMSO: dimethyl sulfoxide
DNA: deoxyribonucleic acid
EC_50_: half maximal effective concentration
ELISA: enzyme-linked immunosorbent assay
*F*_o_ – *F*_o_: F_obs(fragment soaked)_ – F_obs(DMSO blank)_ map
FBDD: fragment-based drug discovery
HIV-1: human immunodeficiency virus-1
IC_50_: half maximal inhibitory concentration
LE: ligand efficiency
NMR: nuclear magnetic resonance
NNIBP: non-nucleoside reverse transcriptase inhibitor binding pocket
NNRTI: non-nucleoside reverse transcriptase inhibitor
NRTI: nucleoside reverse transcriptase inhibitor
PDB: Protein Data Bank
PEG: polyethylene glycol
RNA: ribonucleic acid
RPV: rilpivirine
RT: reverse transcriptase
SBDD: structure-based drug discovery
XCFS: X-ray crystallographic fragment screening

