## Supporting Information for "The Halo Library, a Tool for Rapid Identification of Ligand Binding Sites on Proteins Using Crystallographic Fragment Screening"

Page 1 – **Figure S1** – Fragment binders from Halo Library and the 2013 XCFS campaign bound to RT-rilpivirine (RPV).

Page 1 – **Figure S2** – Knuckles site can be extended into a neighboring pocket.

Page 2 – **Figure S3** – IC_50_ plots for inhibition of the enzymatic activity of HIV-1 RT by Halo Library fragments **HL6**, **HL15**, and **HL20**.

Page 3 – **Table S1** – Chemical names and SMILES strings for fragments in the Halo Library.

Page 5 – **Table S2** – X-ray crystallographic data collection and refinement parameters.

Page 8 – **Table S3** – Electron density maps for Halo Library binders of RT-RPV.

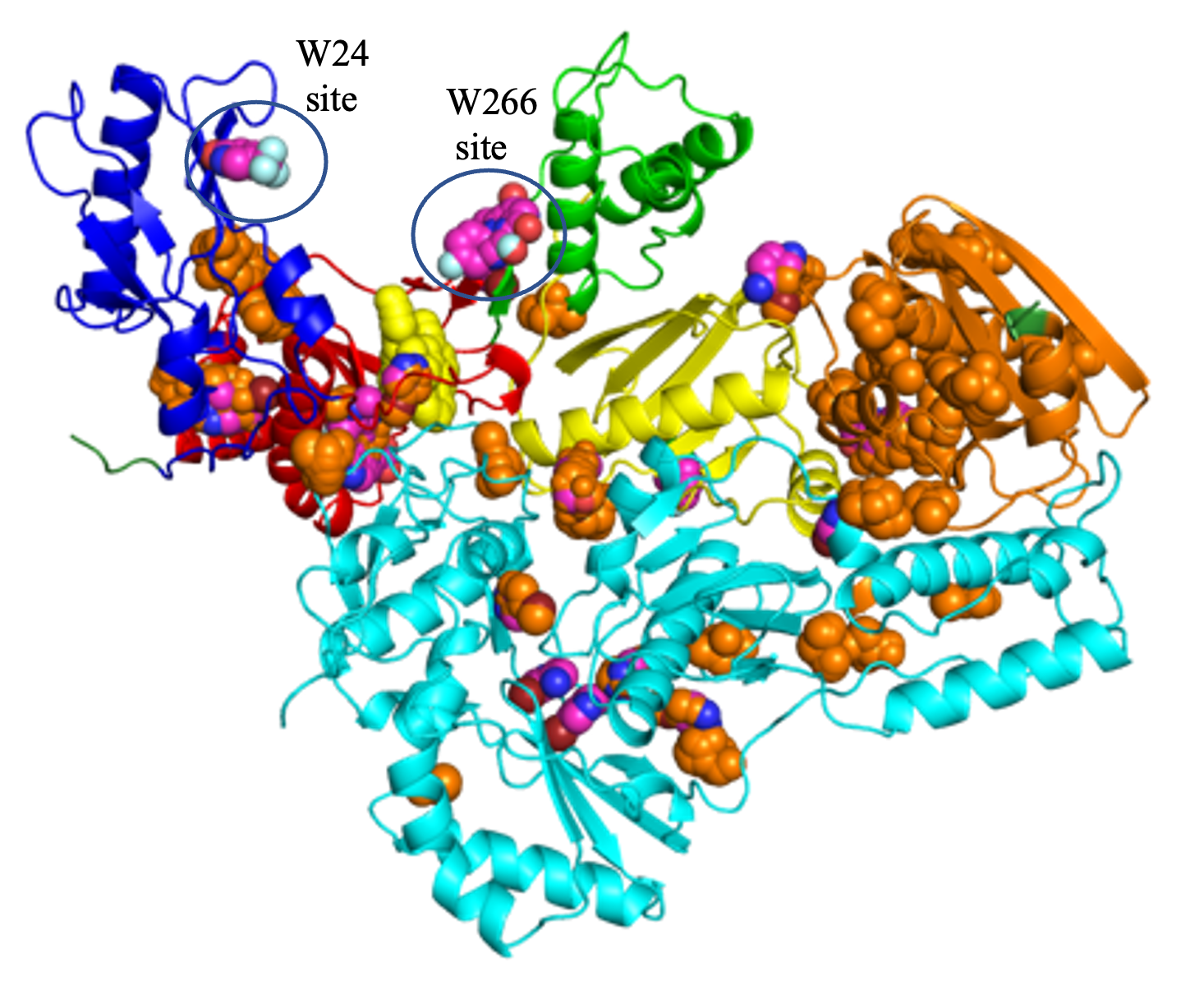

**Figure S1**. Fragment binders from the Halo Library (magenta spheres) and the 2013 XCFS campaign (orange spheres) bound to HIV-1 RT-rilpivirine (RPV).^1^ RT is color coded to show fingers (dark blue), palm (red), thumb (green), connection (yellow), and RNase H (magenta) subdomains of the p66 subunit, and p51 is shown in cyan. RPV is shown as yellow spheres. W24 and W266 sites, which were discovered in the Halo campaign but not in the 2013 campaign, are indicated.

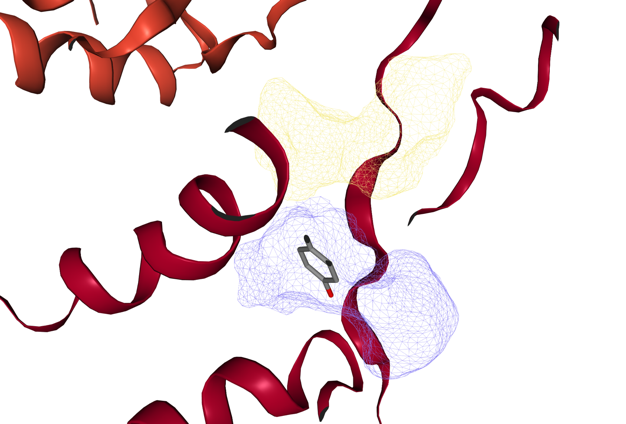

**Figure S2.** Knuckles site (purple mesh) can be extended into a neighboring pocket (yellow mesh) as found in DoGSiteScorer and discussed in text.

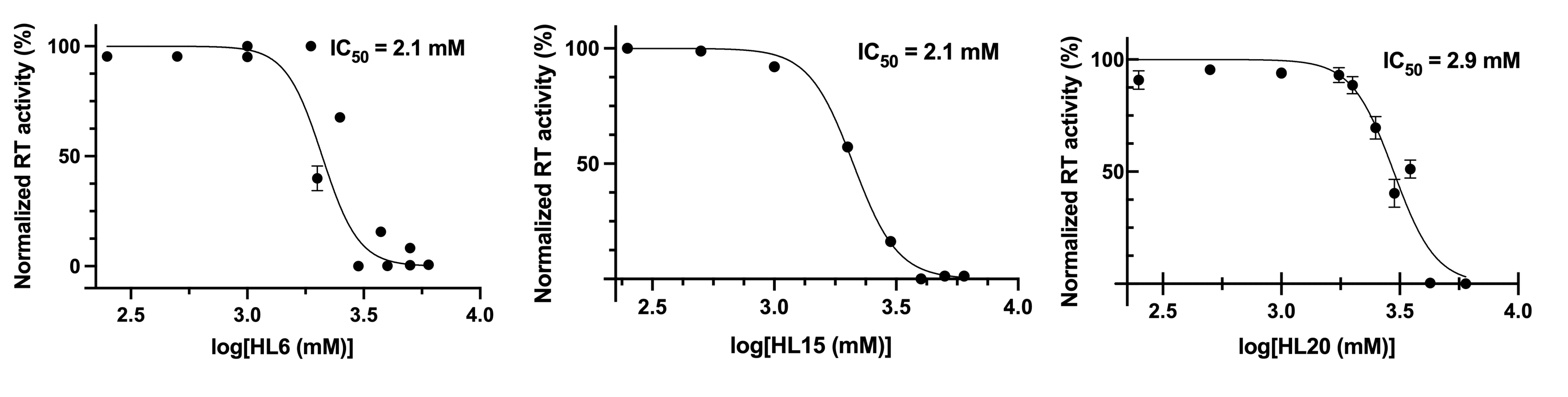
**Figure S3.** IC_50_ plots for inhibition of the enzymatic activity of HIV-1 RT by Halo Library fragments **HL6**, **HL15**, and **HL20**, with values indicated on the top right of each plot.

1.	Bauman, J. D.; Patel, D.; Dharia, C.; Fromer, M. W.; Ahmed, S.; Frenkel, Y.; Vijayan, R. S. K.; Eck, J. T.; Ho, W. C.; Das, K.; Shatkin, A. J.; Arnold, E., Detecting Allosteric Sites of HIV-1 Reverse Transcriptase by X-ray Crystallographic Fragment Screening. *J. Med. Chem.* **2013,** *56* (7), 2738-2746.

**Table S1. Chemical names and SMILES strings for fragments in the Halo Library.**

| **S. NO** | **Name** | **SMILES** |
| --- | --- | --- |
| **HL1** | bromobenzene | BrC1=CC=CC=C1 |
| **HL2** | 3-amino-4-bromopyrazole | BrC1=CNN=C1N |
| **HL3** | 3-bromo thiophene | BrC1=CSC=C1 |
| **HL4** | 5-bromo-2-furoic acid | O=C(O)C1=CC=C(Br)O1 |
| **HL5** | 2-bromo anisole | COC1=CC=CC=C1Br |
| **HL6** | 4-amino-3-bromopyridine | BrC1=C(N)C=CN=C1 |
| **HL7** | 3-bromoimidazo[1,2-a]pyridine | BrC1=CN=C2C=CC=CN21 |
| **HL8** | 2-bromoacetophenone | O=C(C1=CC=CC=C1)CBr |
| **HL9** | 3-bromobenzylaminehydrochloride | NCC1=CC=CC(Br)=C1 |
| **HL10** | 4-bromo-3-methylpyrazole | CC1=NNC=C1Br |
| **HL11** | 2-bromopyrimidine | BrC1=NC=CC=N1 |
| **HL12** | 3-bromo-N-methylaniline | CNC1=CC=CC(Br)=C1 |
| **HL13** | 5-amino-2-bromopyridine | BrC1=NC=C(N)C=C1 |
| **HL14** | 5-bromopyrimidine | BrC1=CN=CN=C1 |
| **HL15** | 2-chloro-6-fluorophenethylamine | NCCC1=C(F)C=CC=C1Cl |
| **HL16** | 2-(4-fluorophenyl)-2-propanol | CC(O)(C1=CC=C(F)C=C1)C |
| **HL17** | 1,2- dimethoxy-4-fluorobenzene | FC1=CC=C(OC)C(OC)=C1 |
| **HL18** | 4-fluoroaniline | NC1=CC=C(F)C=C1 |
| **HL19** | 4-fluorophenylurea | O=C(N)NC1=CC=C(F)C=C1 |
| **HL20** | 5-fluoroindole-2-carboxylic acid | O=C(C(N1)=CC2=C1C=CC(F)=C2)O |
| **HL21** | 1-bromo-4-fluoro-2-iodobenzene | IC1=CC(F)=CC=C1Br |
| **HL22** | 1-bromo-2,3,4-trifluorobenzene | FC1=CC=C(Br)C(F)=C1F |
| **HL23** | 2-amino-6-fluorobenzonitrile | N#CC1=C(F)C=CC=C1N |
| **HL24** | 4-(trifluoromethyl) cyclohexan-1-ol | OC1CCC(C(F)(F)F)CC1 |
| **HL25** | 1-methyl-5-(trifluoromethyl)pyrazol-3-ol | OC1=NN(C)C(C(F)(F)F)=C1 |
| **HL26** | 4-(3-fluorophenyl)-2-methyl-1,3-thiazole | CC1=NC(C2=CC=CC(F)=C2)=CS1 |
| **HL27** | 5-(trifluoromethyl)pyridin-2-ol | OC1=NC=C(C(F)(F)F)C=C1 |
| **HL28** | 5-fluoro-4-sulfanylidene-1,3-dihydropyrimidin-2-one | O=C1NC(C(F)=CN1)=S |
| **HL29** | 5-fluoroquinazolin-4-ol | OC1=C2C(F)=CC=CC2=NC=N1 |
| **HL30** | 4-[(trifluoromethyl)]sulfanylbenzamide | O=C(N)C1=CC=C(SC(F)(F)F)C=C1 |
| **HL31** | 4- bromophenol | OC1=CC=C(Br)C=C1 |
| **HL32** | 2-bromobenzoic acid | O=C(O)C1=CC=CC=C1Br |
| **HL33** | 2- amino-5-fluorobenzoic acid | O=C(O)C1=CC(F)=CC=C1N |
| **HL34** | [1-(4-fluorophenyl)-5-methyl-1H-pyrazol-4-yl]methanol | OCC1=C(C)N(C2=CC=C(F)C=C2)N=C1 |
| **HL35** | 4-fluorophenol | OC1=CC=C(F)C=C1 |
| **HL36** | iodobenzene | IC1=CC=CC=C1 |
| **HL37** | 4-bromo-1H-imidazole | BrC1=CNC=N1 |
| **HL38** | 1-(4-bromophenyl) ethanone | CC(C1=CC=C(Br)C=C1)=O |
| **HL39** | 2-bromobenzoic acid | O=C(O)C1=CC=CC=C1Br |
| **HL40** | 4-(2-aminoethyl) benzene sulfonyl fluoride | O=S(C1=CC=C(CCN)C=C1)(F)=O |
| **HL41** | 5-fluorouracil | O=C(N1)NC=C(F)C1=O |
| **HL42** | 5-fluoro-1H-indole | FC1=CC2=C(NC=C2)C=C1 |
| **HL43** | 1-N-methyl-4-(trifluoromethyl)benzene-1,2-diamine | NC1=CC(C(F)(F)F)=CC=C1NC |
| **HL44** | 4-bromopyrazole | BrC1=CNN=C1 |
| **HL45** | 4-fluoro-1H-pyrazole | FC1=CNN=C1 |
| **HL46** | 4-iodopyrazole | IC1=CNN=C1 |

**Table S2. X-ray crystallographic data collection and refinement parameters for Halo Library fragment binders of RT-RPV crystals.**

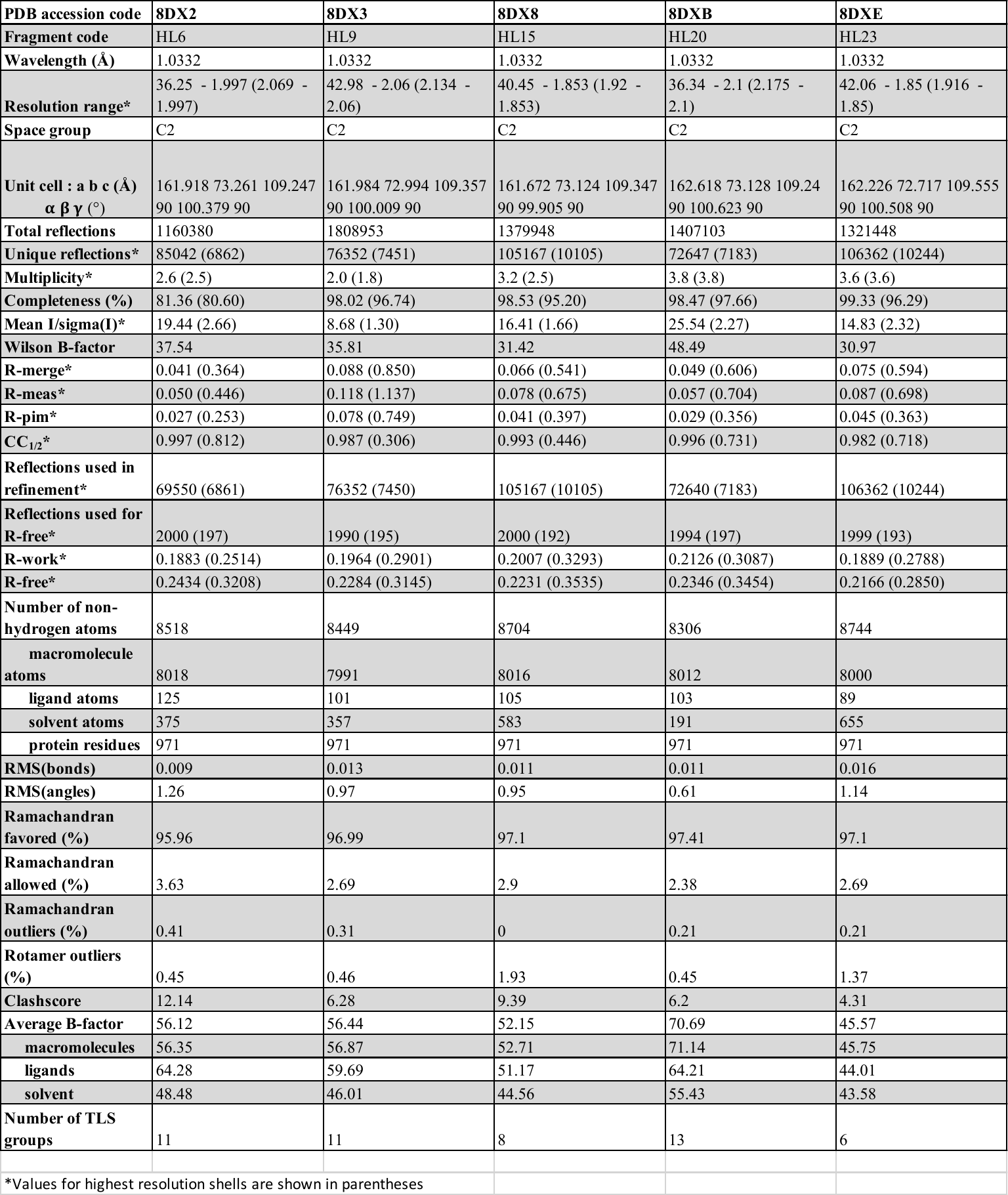

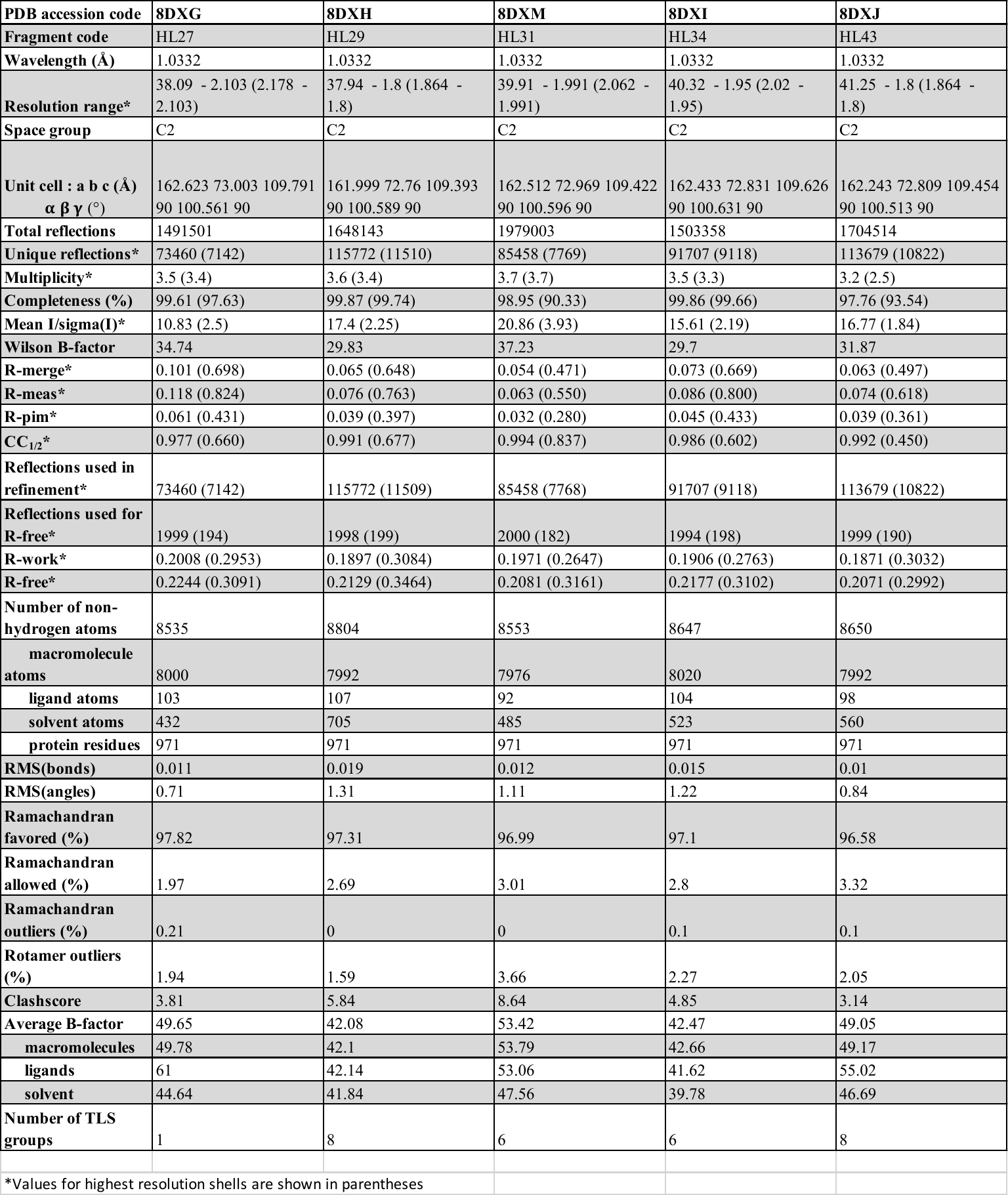

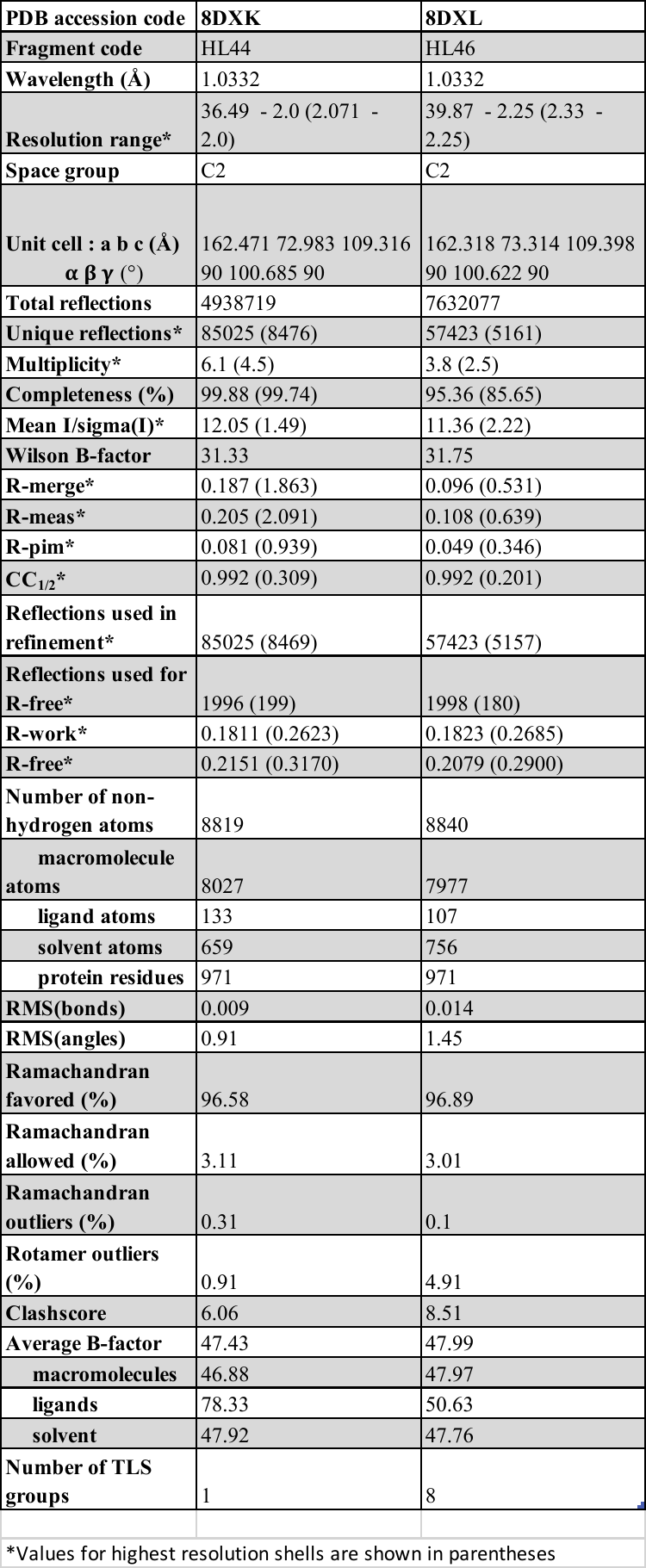

**Table S3. Electron density maps for Halo Library binders of RT-RPV discussed in text. *F*_o_ – *F*_o_ difference maps (dark green, σ = 3.0), 2*F*_o_ ­– *F*_c_ maps (blue, σ = 1.0), and Polder OMIT maps (light green, σ = 3.0) are shown. Some maps are shown at different σ values, which are indicated on the bottom right of each cell.**

| **Fragment (Binding site)** | ***PDB ID*** | ***F*_o_ – *F*_o_ difference map** | **2*F*_o_ ­– *F*_c_ map** | **Polder OMIT map** |
| --- | --- | --- | --- | --- |
| **HL6** (NNRTI adjacent) | 8DX2 | 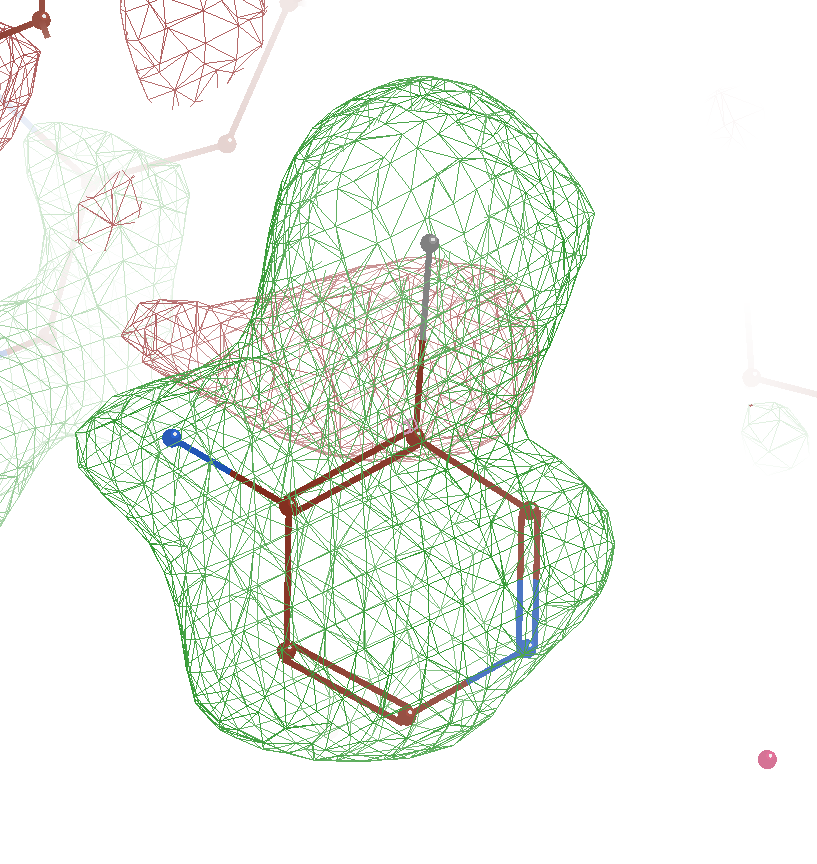 | 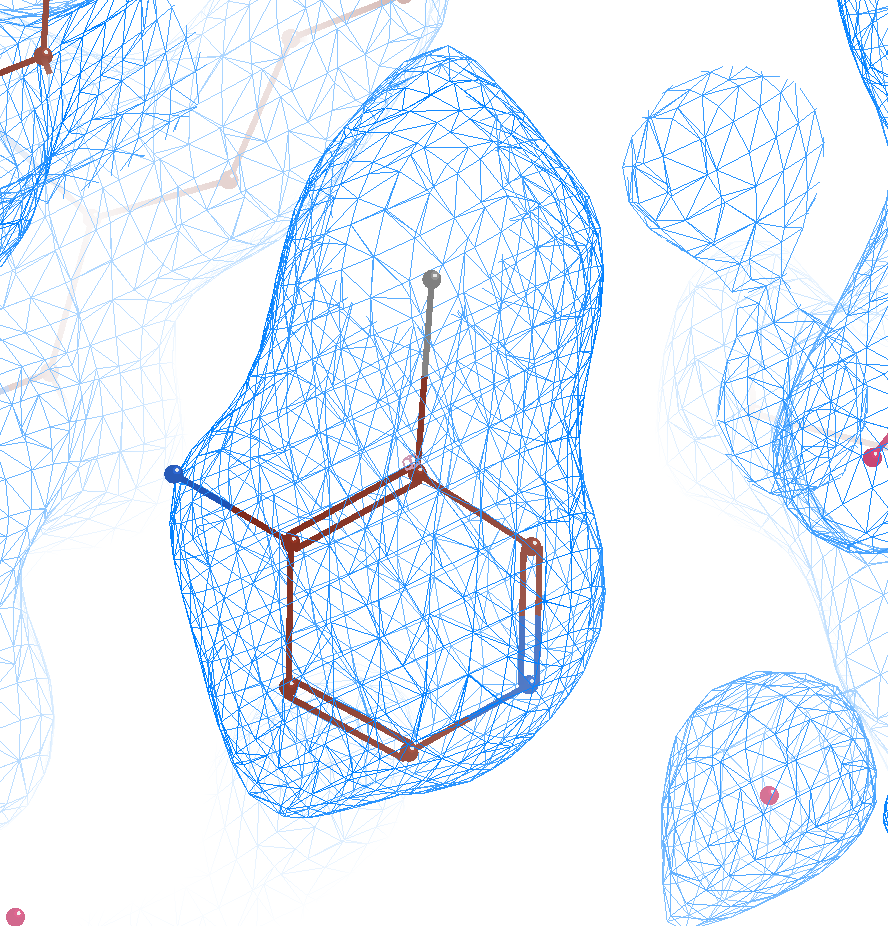 | 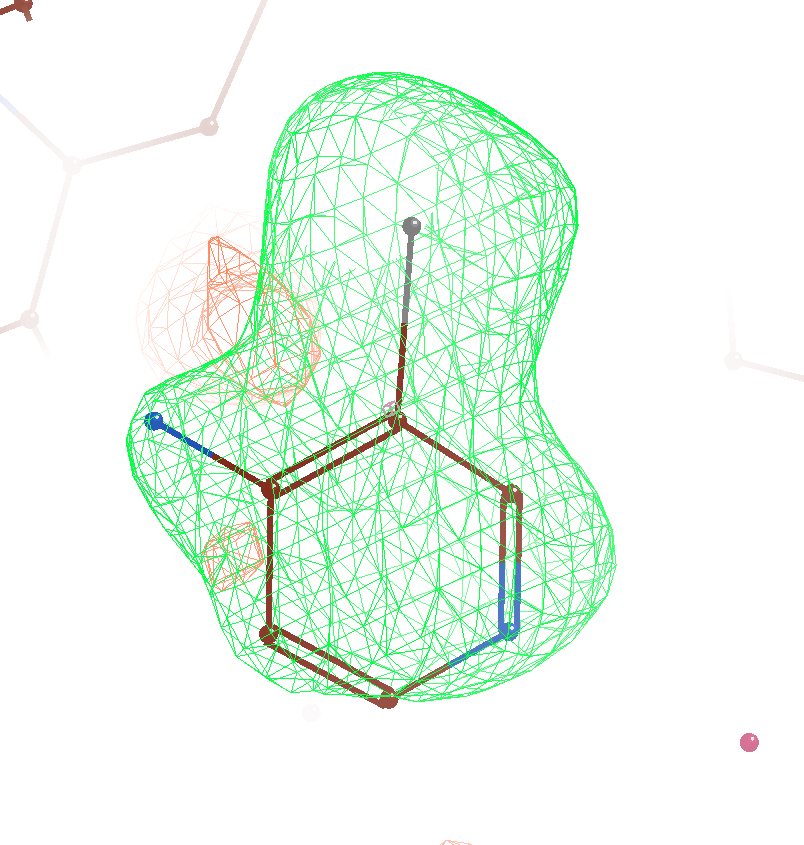 |
| **HL20** (NNRTI adjacent) | 8DXB | 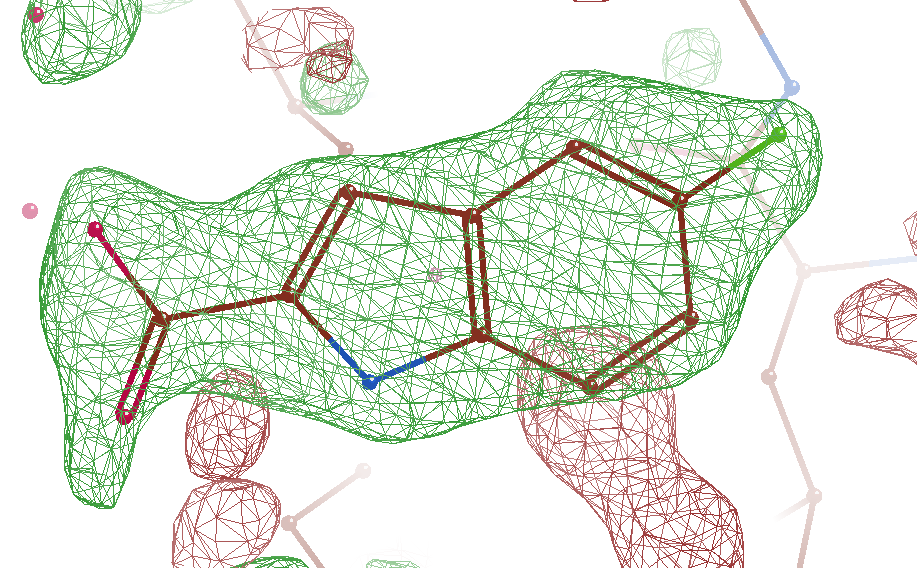 | 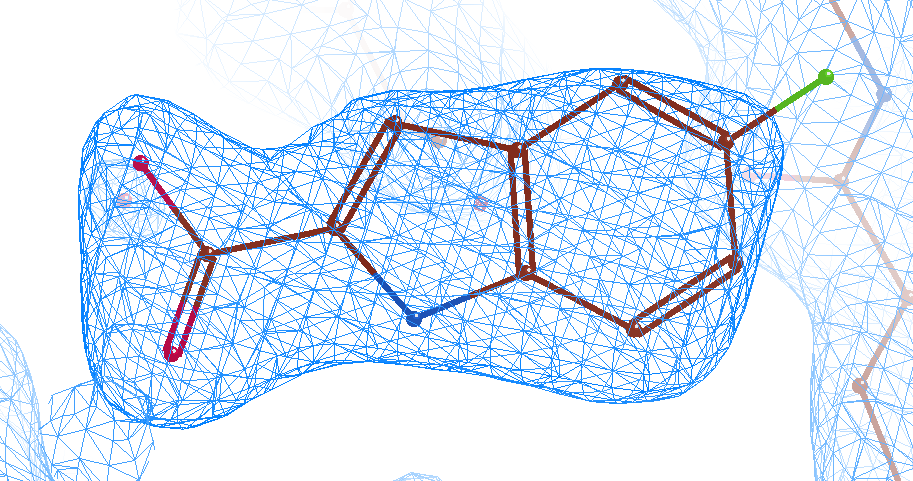 | 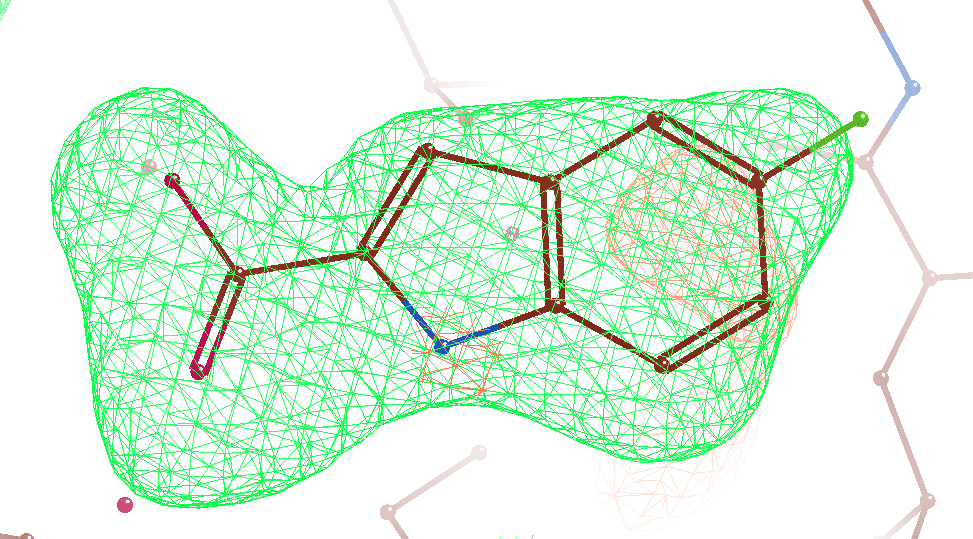 |
| **HL23** (NNRTI adjacent) | 8DXE | 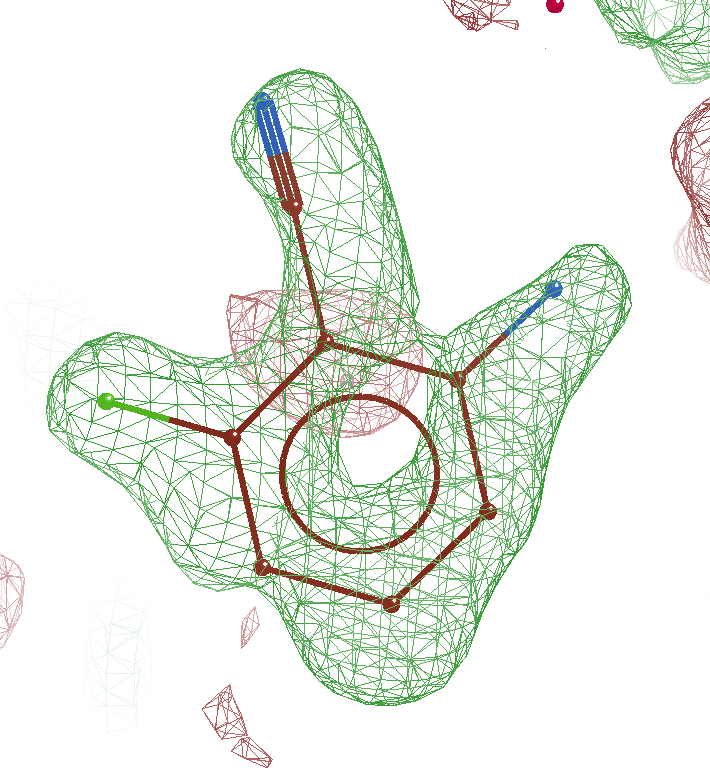 | 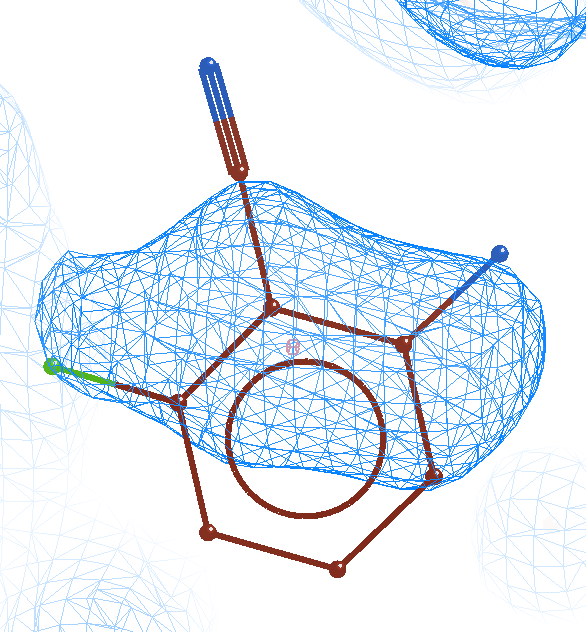 | 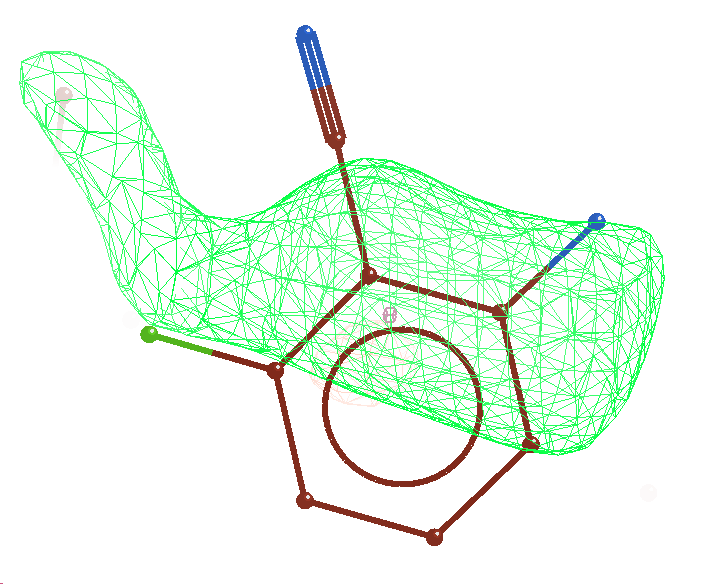 |
| **HL43** (NNRTI adjacent) | 8DXJ | 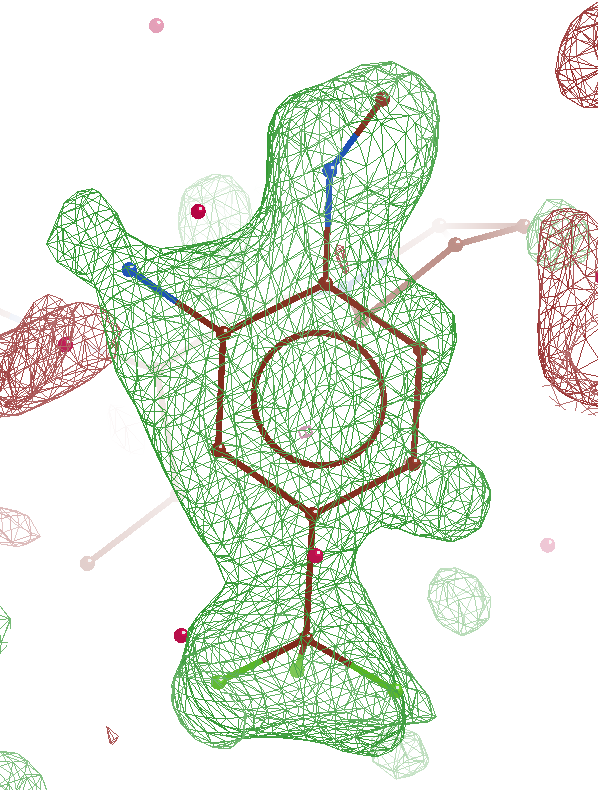 | 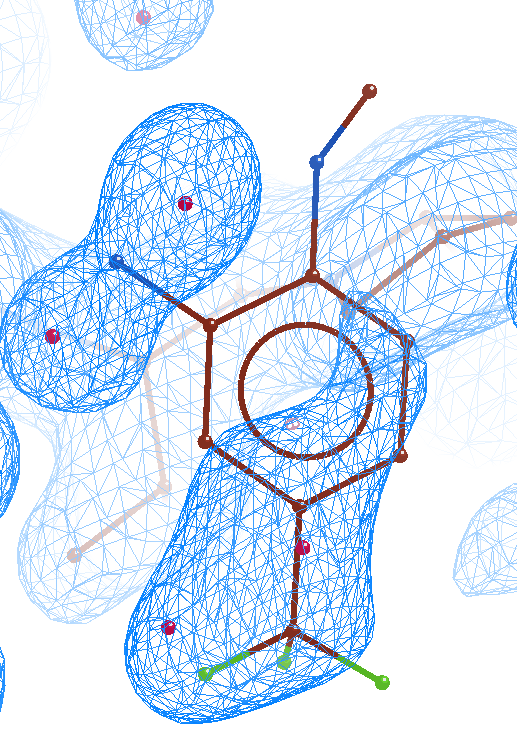 | 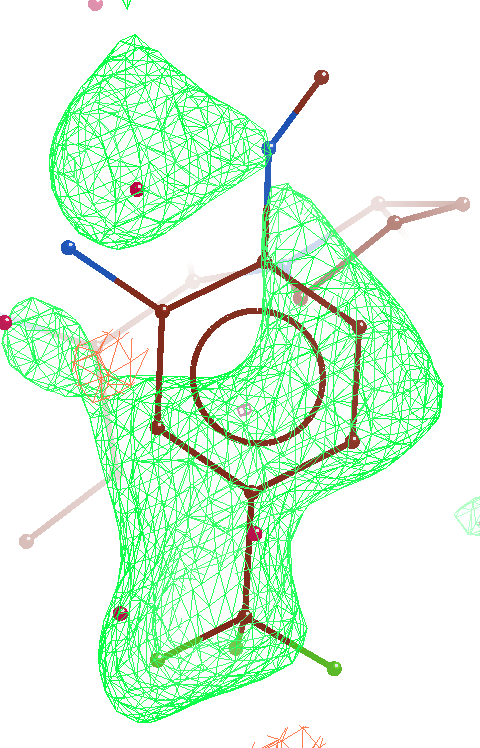 |
| **HL44** (NNRTI adjacent) | 8DXK | 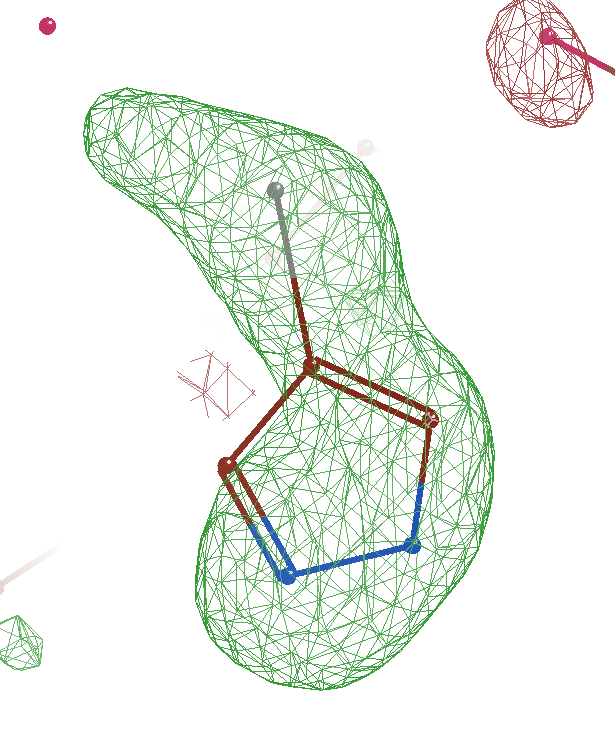 | 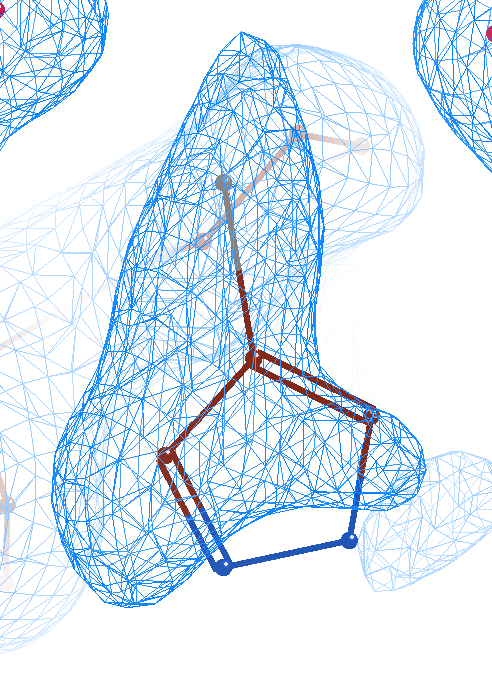 | 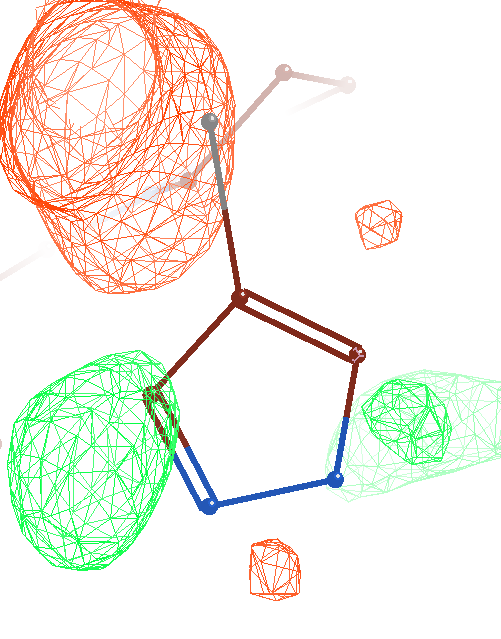 |
| **HL46** (NNRTI adjacent) | 8DXL | 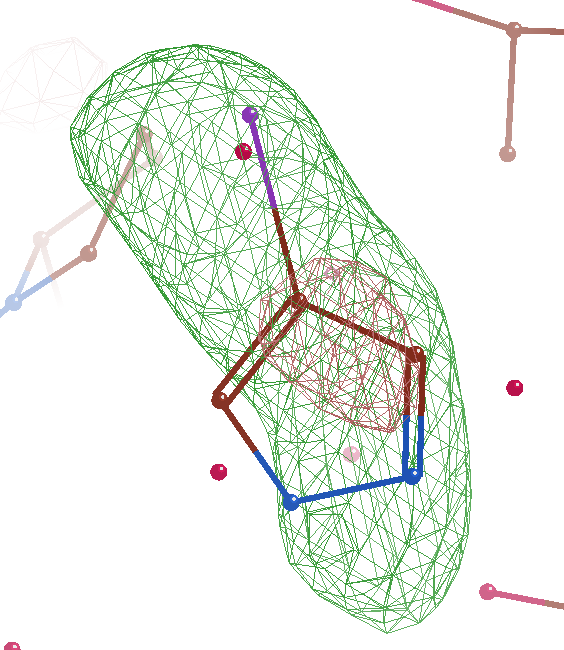 | 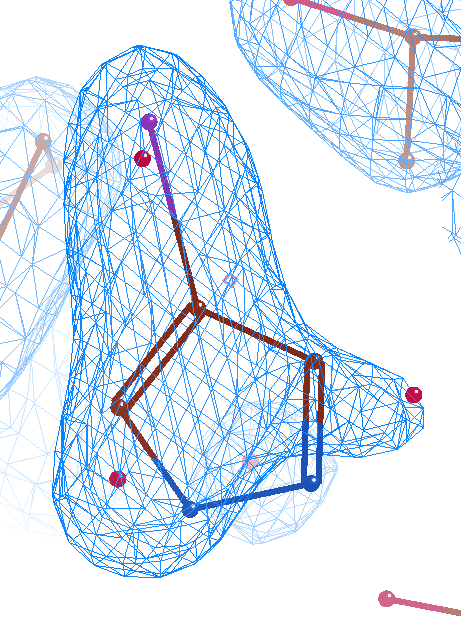 | 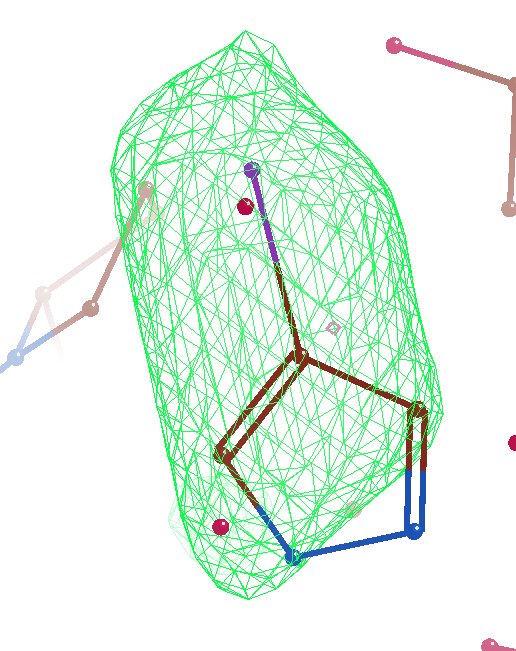3.8 |
| **HL27** (W24) | 8DXG | 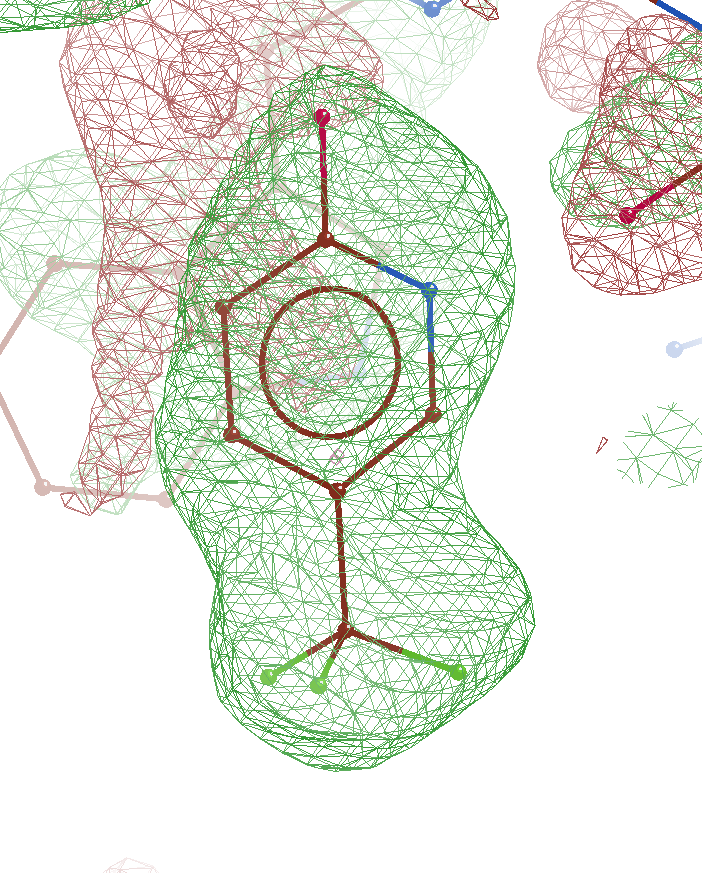 | 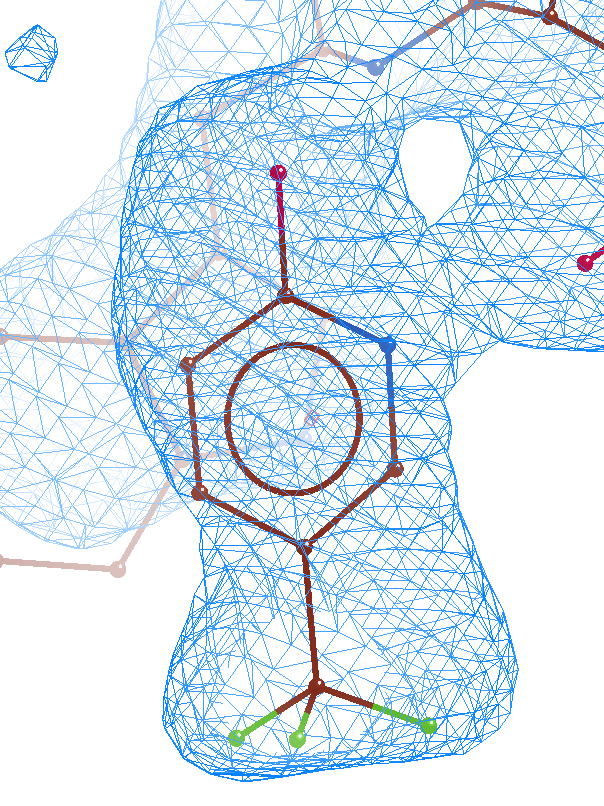 | 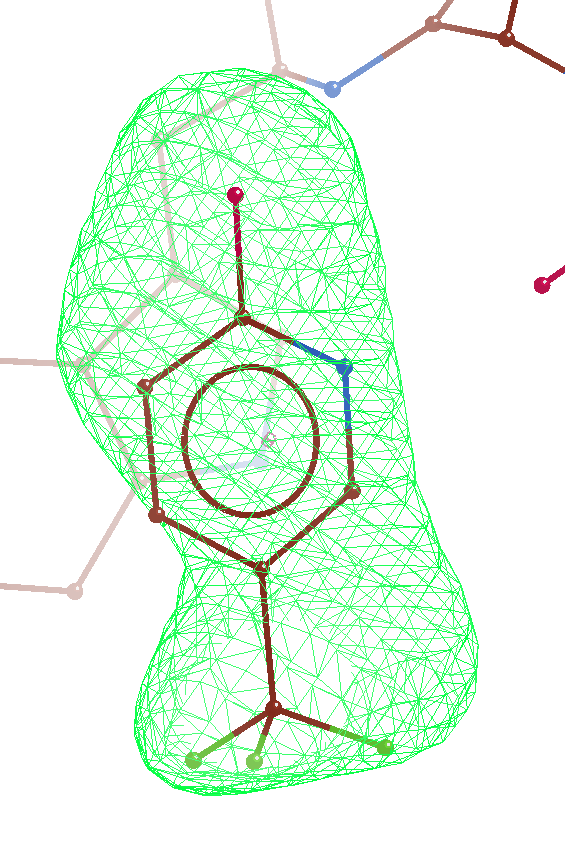 |
| **HL15** (415) | 8DX8 | 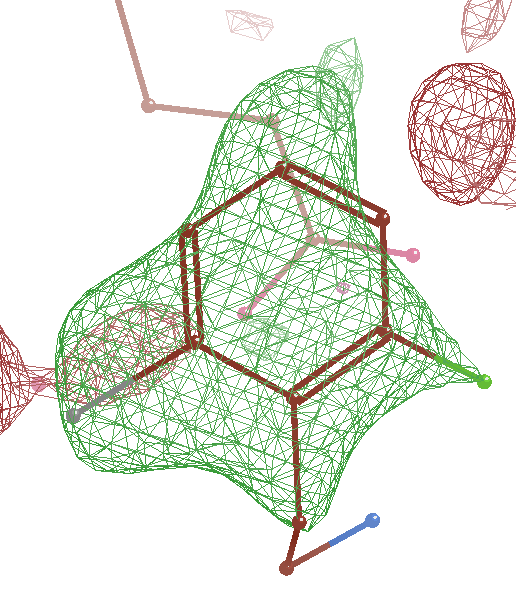 | 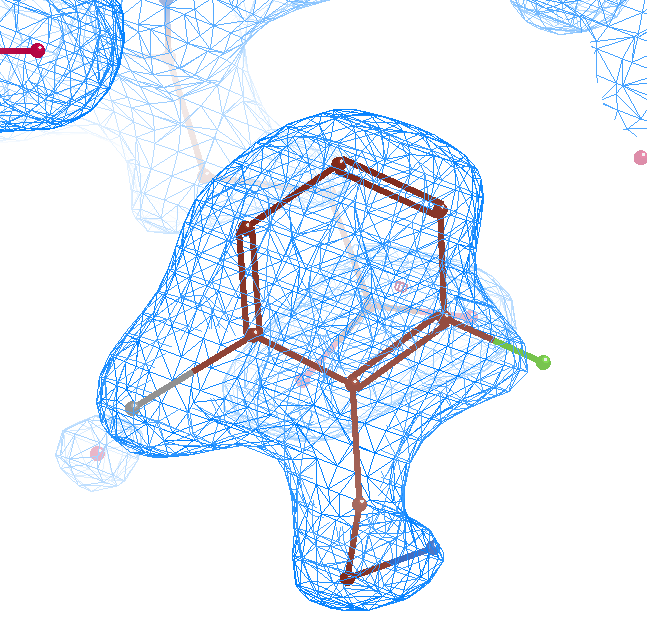 | 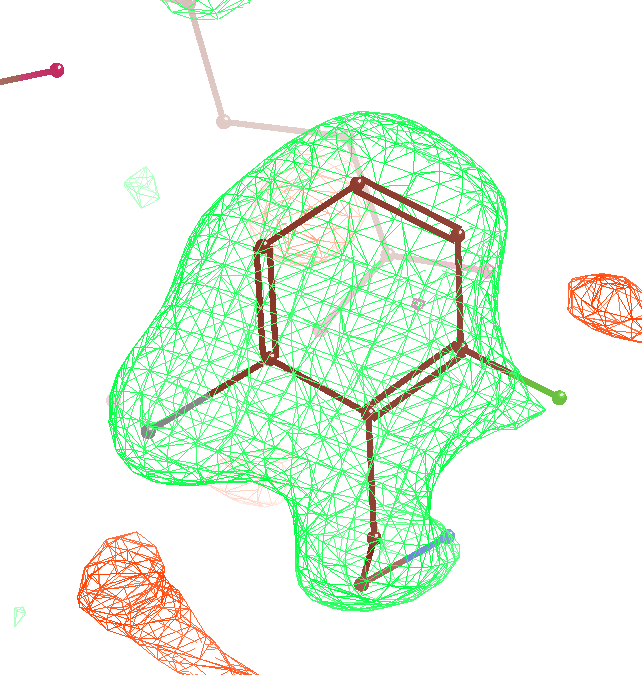 |
| **HL29** (W266) | 8DXH | 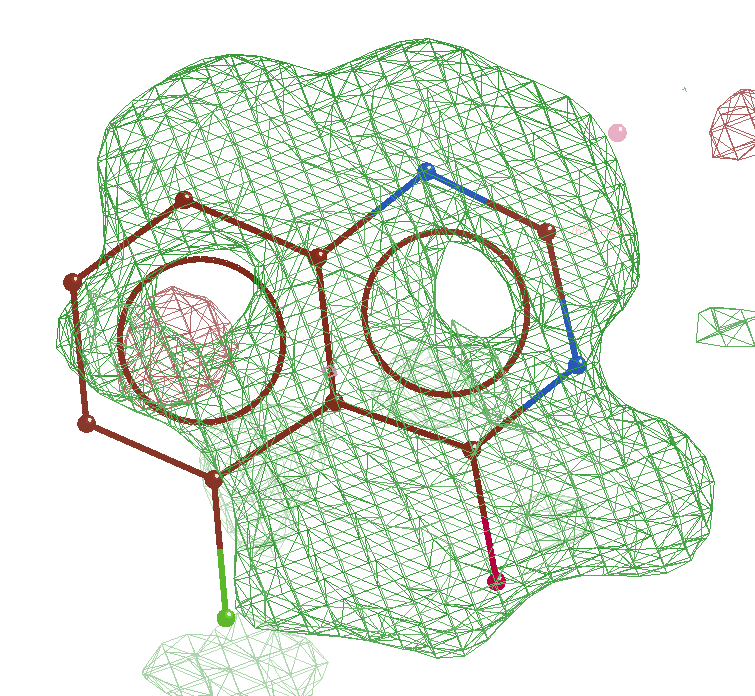 | 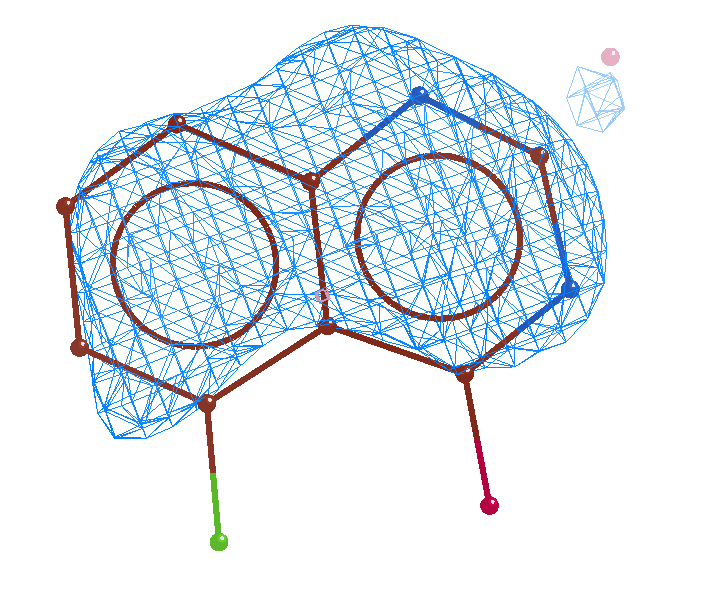 | 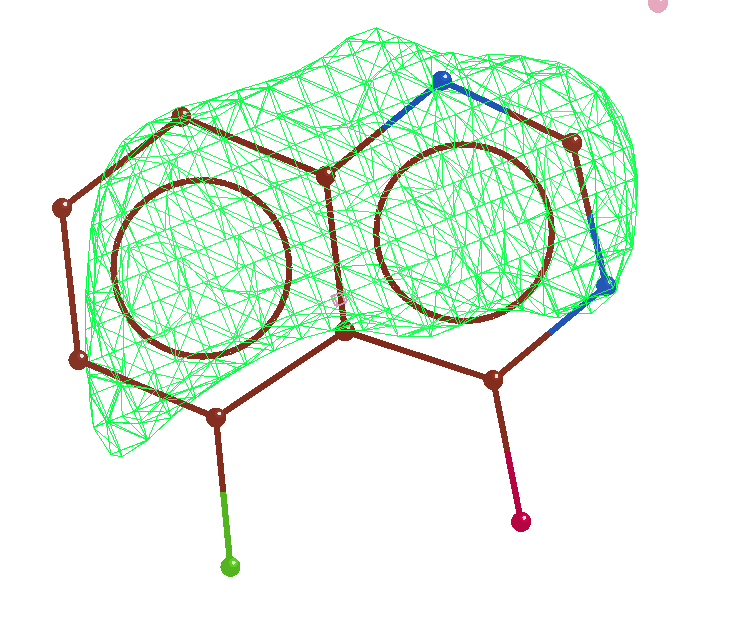 |
| **HL34** (W266) | 8DXI | 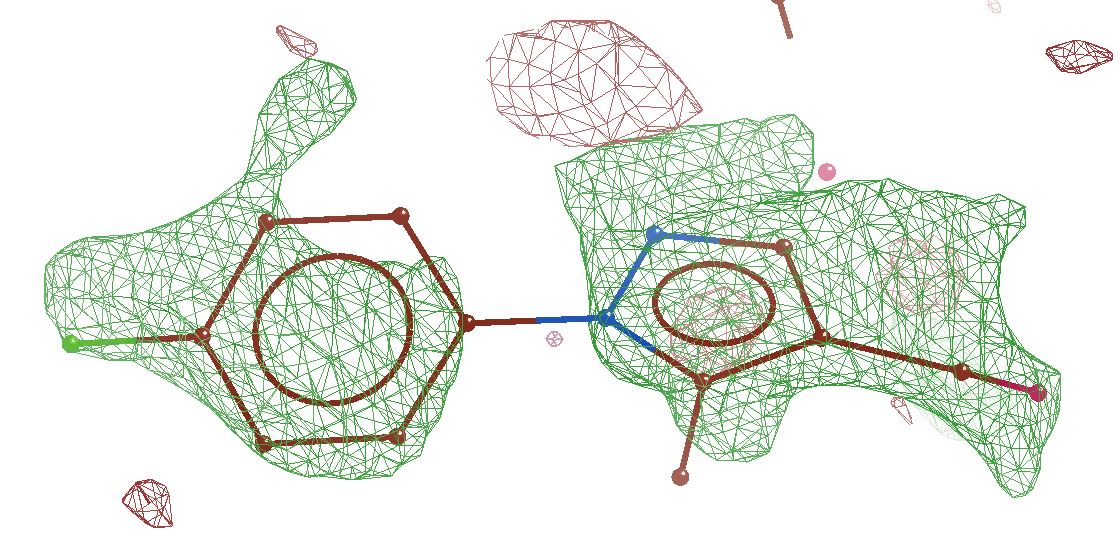 | 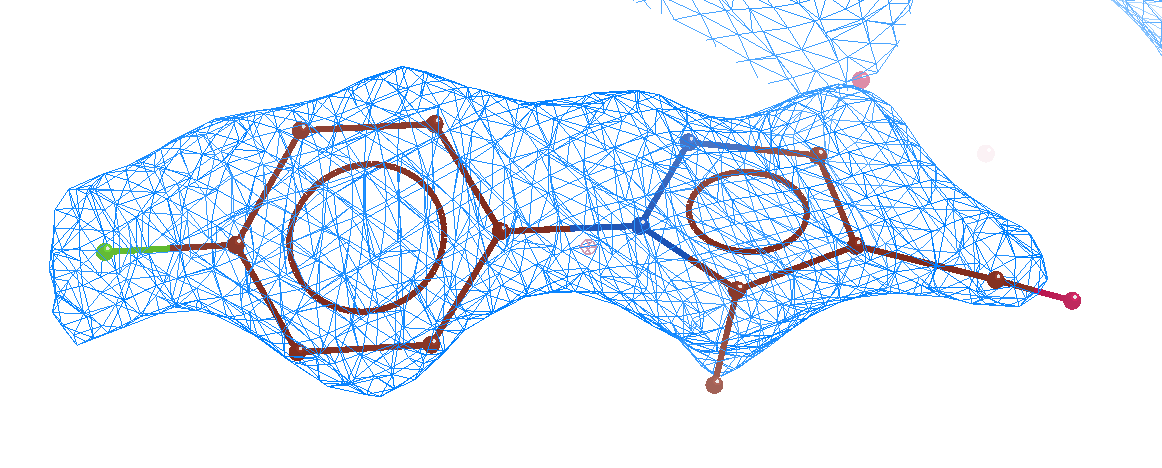 | 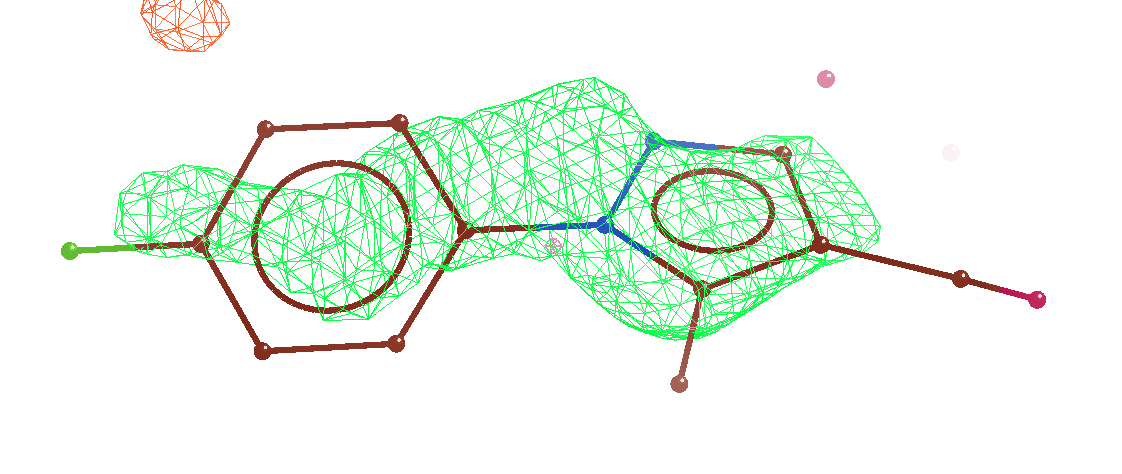 |
| **HL31** (Knuckles) | 8DXM | 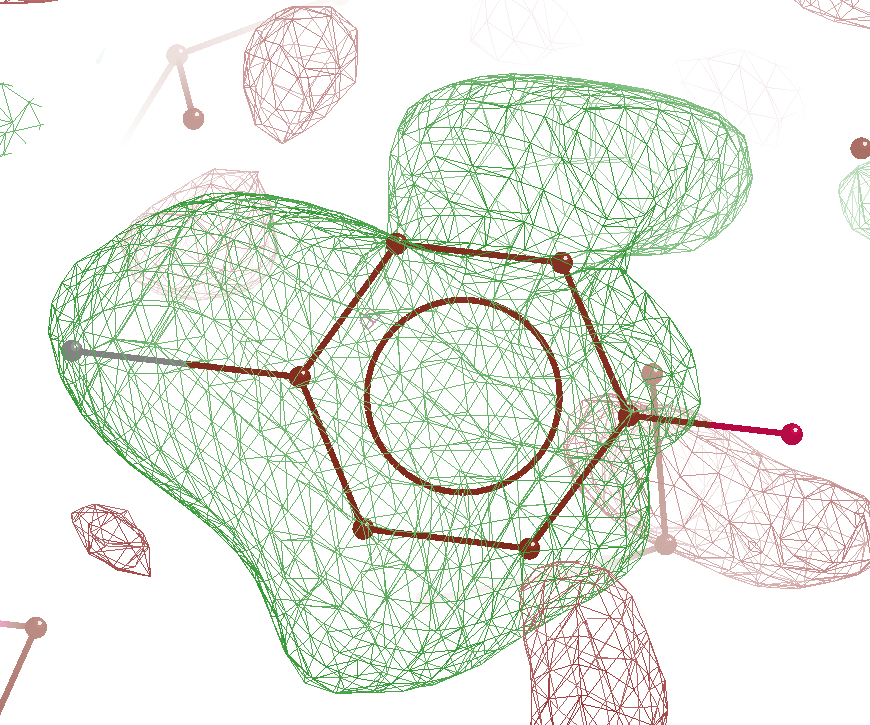 | 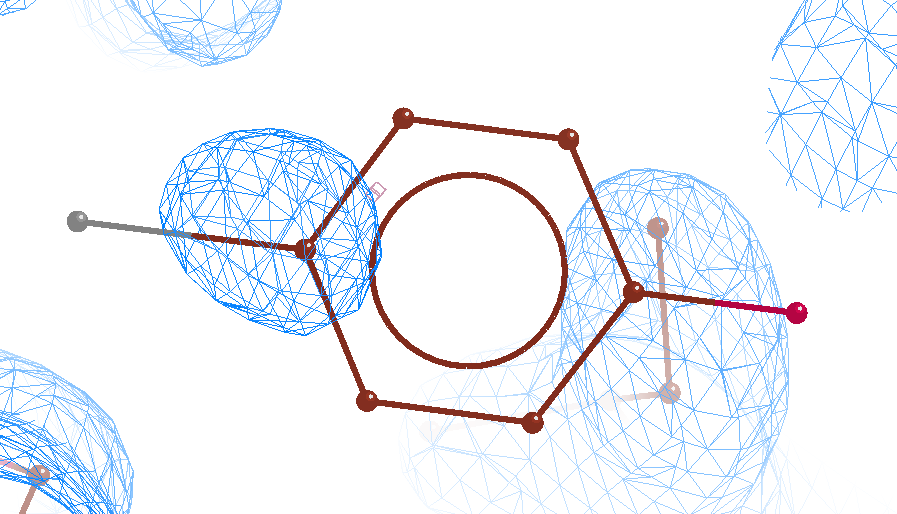 | 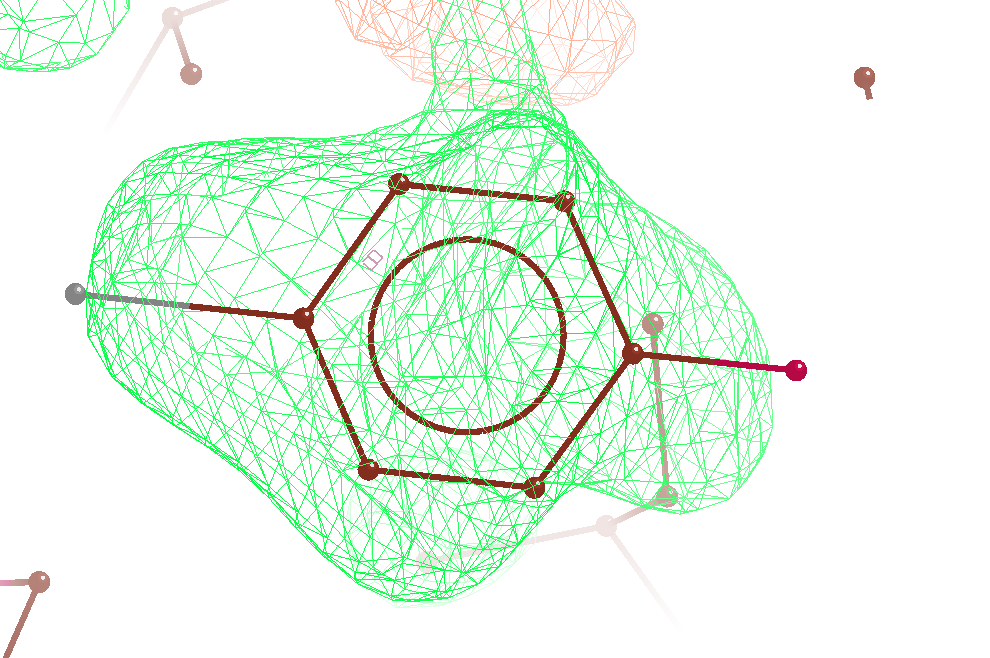 |
| **HL46** (Knuckles) | 8DXL | 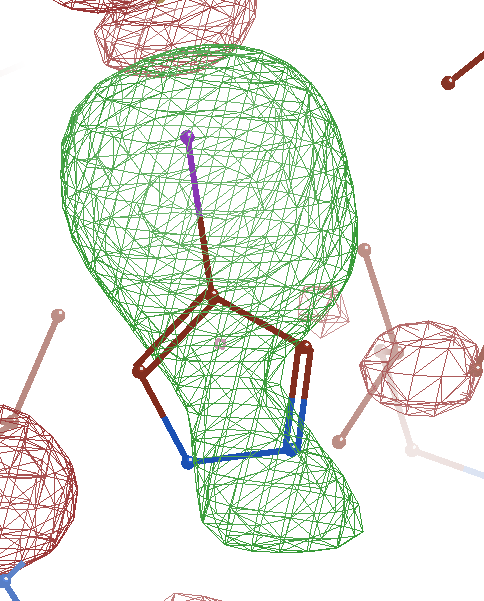 | 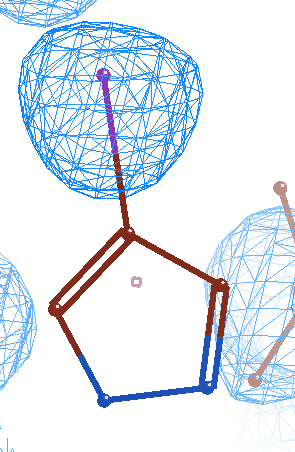 | 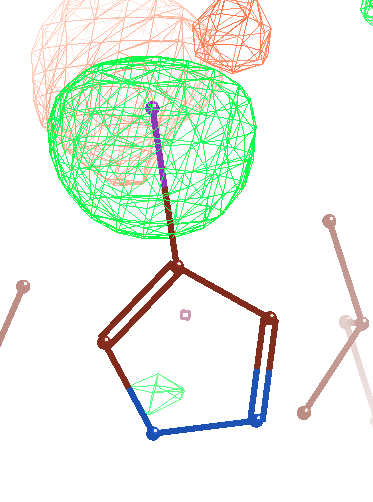 |
